# CESA7 and microtubules pattern complex secondary cell walls in explosive fruit

**DOI:** 10.64898/2026.01.25.701578

**Authors:** Ryan C. Eng, Aurélia Emonet, Ulla Neumann, Markus Pauly, Angela Hay

## Abstract

Secondary cell walls (SCW) constitute the most abundant form of renewable plant biomass and are major sinks for atmospheric carbon. Their highly ordered patterns underpin specialized cell functions. In *Cardamine hirsuta*, the geometry of a polarly localized SCW in fruit endocarp *b* (end*b*) cells determines the mechanics of explosive seed dispersal. Yet, the genetic control of SCW synthesis and patterning in these specialized cells remains poorly understood. Here we show that *CELLULOSE SYNTHASE 7* (*CESA7*) is required to synthesize SCW cellulose in end*b* cells. While lignin and xylan deposition occurs independently of cellulose patterning in *cesa7* end*b* SCWs, the final geometry and layered organization of wild-type end*b* SCWs depend on *CESA7*. Cellulose serves as a scaffold for the organized assembly of SCW polymers, thereby maintaining the precise SCW patterns observed in end*b* cells of fruits and metaxylem cells in roots. Cortical microtubules guide the patterned deposition of cellulose, lignin and xylan in end*b* cells, creating SCW-depleted domains along cell edges that produce the specific hinged SCW geometry. Disrupting microtubules abolished this pattern and prevented explosive coiling of the fruit valves. Our findings show that microtubules and *CESA7* shape the form and function of end*b* SCWs in exploding seed pods.

## Introduction

Explosive seed dispersal is employed by the common weed *Cardamine hirsuta* (hairy bittercress) to fling its seeds far from the plant and effectively colonize disturbed land. This trait distinguishes species in the *Cardamine* genus from the related model plant Arabidopsis (*Arabidopsis thaliana*), which has non-explosive seed pods (Emonet and Hay, 2024). In *C. hirsuta*, the two valves of an exploding seed pod coil in an ultrafast movement, launching seeds on ballistic trajectories at speeds faster than 10 m/s (Hofhuis et al., 2016). Key innovations for this trait to evolve included mechanisms to store and rapidly release elastic energy (Cullen and Hay, 2024). The tension that generates elastic energy is produced in growing fruit by differential contraction of valve tissues (Mosca et al., 2024). Explosive release of this tension is controlled at the cellular scale by polar deposition of a lignified secondary cell wall (SCW) in endocarp *b* (end*b*) cells of the fruit valve (Hofhuis et al., 2016). Switches in end*b* SCW patterning between uniform and polar underlie trait transitions between non-explosive and explosive seed dispersal both within and between closely related species (Hofhuis et al., 2016; Hofhuis and Hay, 2017; Emonet et al., 2024). Thus, to understand the development and evolution of explosive seed dispersal, it is important to identify genetic regulators of end*b* SCW synthesis and patterning.

Explosive seed dispersal in *C. hirsuta* depends on SCW formation in end*b* cells, as mutants lacking this layer fail to explode (Hofhuis et al., 2016). The hinged geometry of end*b* SCWs enables fruit valves to coil rapidly, like a toy slap bracelet, releasing stored elastic energy (Hofhuis et al., 2016). By contrast, non-explosive seed dispersal in Arabidopsis relies on SCW formation in the dehiscence zone rather than in the end*b* cell layer (Mitsuda and Ohme-Takagi, 2008), highlighting species-specific functions of SCWs in seed dispersal.

To identify genes required for the lignified end*b* SCW in *C. hirsuta*, a genetic screen was previously conducted for mutants with less lignified fruit valves. The transcription factor SQUAMOSA PROMOTER-BINDING PROTEIN-LIKE 7 (SPL7) was identified as a regulator of copper homeostasis, required for robust lignification of end*b* SCWs (Perez-Anton et al., 2022). Three multicopper laccases, LAC4, 11, 17, were found to precisely colocalize with the lignified end*b* SCW, and loss of all three genes resulted in complete loss of lignin in *lac4 11 17* end*b* SCWs (Perez-Anton et al., 2022). Laccases are secreted enzymes that function in the cell wall to activate monolignols into radicals by oxidation. The lignin polymer then forms by nonenzymatic random coupling of these activated monolignols (Dixon and Barros, 2019). Therefore, the specific localization of LAC4, 11, 17, embedded in the polysaccharide cell wall matrix, determines precisely where lignin is deposited in end*b* SCWs. *In situ* laccase activity assays bridged these findings by showing that these three oxidative enzymes depend on the SPL7 pathway to provide sufficient copper for their catalytic activity in end*b* SCWs (Perez-Anton et al., 2022). Although the end*b* SCW of *spl7* and *lac4 11 17* mutants lacked lignin, the layered organization and hinged pattern of the SCW remained similar to wild type (Perez-Anton et al., 2022). Cellulose staining of the non-lignified end*b* SCW of these mutants (Perez-Anton et al., 2022) suggested that the templated synthesis of cellulose may be critical for generating the distinctive SCW pattern in *C. hirsuta* end*b* cells.

Cellulose microfibrils, composed of linear β-1,4-linked glucose polymers, are synthesized at the plasma membrane by cellulose synthase complexes (CSCs) (McFarlane et al., 2014). Each CSC is a hexamer of cellulose synthase (CESA) heterotrimers, producing 18 linear cellulose chains (Kubicki et al., 2018; Purushotham et al., 2020). These chains crystallize at the outer side of the membrane into a stiff microfibril, which might further coalesce into higher order microfibril structures in SCWs (Cosgrove et al., 2024). A cryo-EM structure of a CESA trimer showed that a large channel forms a path for cellulose chains through the membrane-embedded complex (Purushotham et al., 2020). In this way, CESA proteins, which belong to the glycosyltransferase-2 superfamily (Pear et al., 1996), combine two functions: synthesizing cellulose and secreting the polymer through a transmembrane channel formed by its membrane-spanning segment (Pedersen et al., 2023). In Arabidopsis, 10 CESA isoforms exist (Richmond and Somerville, 2000), with CESA1, CESA3 and one of CESA2/5/6/9 required for cellulose synthesis in primary cell walls (Desprez et al., 2007; Persson et al., 2007), and CESA4, CESA7 and CESA8 required for cellulose synthesis in SCWs (Taylor et al., 2003). CESA composition is exchanged during the transition from primary to SCW formation by the turnover of primary wall CSCs and the delivery of SCW CSCs to the plasma membrane (Watanabe et al., 2018). All three SCW CESA subunits are required in order to assemble a CSC for SCW cellulose synthesis, therefore strong loss-of-function alleles of any of these three genes resulted in little or no cellulose in Arabidopsis SCWs (Ha et al., 2002; Taylor et al., 2003).

The formation of thick, cellulose-rich SCWs in specific patterns is associated with distinct CSC properties. Visualizing CESA7-containing CSCs in Arabidopsis epidermal cells by using an ectopic xylem induction system, showed that cellulose was synthesized faster in SCWs, compared to primary walls, due to increased velocity and density of CSCs (Watanabe et al., 2015). Cortical microtubule arrays act to guide the trajectories of CSCs as they move through the plasma membrane, propelled by the extrusion of cellulose microfibrils into the wall (Paredez et al., 2006). Subsequent CSC trajectories can also be guided by the existing cell wall structure, independent of microtubules (Schneider et al., 2017; Chan and Coen, 2020). During SCW formation in protoxylem cells, cortical microtubules are rearranged into banded patterns and coordinate the tracking of CSCs in these banded SCW domains (Watanabe et al., 2015; Schneider et al., 2021). This patterning was lost when microtubules were disrupted by depolymerization, resulting in evenly dispersed CSCs in the plasma membrane (Watanabe et al., 2015). Regulated microtubule depolymerization plays an important role in SCW patterning by creating microtubule-depleted domains adjacent to the plasma membrane that exclude SCW synthesis (Oda et al., 2010; Oda and Fukuda, 2013; Higa et al., 2024). These SCW-free domains form pits in metaxylem cells and the gaps between SCW bands in protoxylem cells (Xu et al., 2022). Therefore, SCW patterning relies on templated cellulose synthesis that is spatially guided by microtubules.

Hemicelluloses, particularly xylan and glucomannan, are the other major constituents of the polysaccharide matrix in SCWs. Both cellulose and xylan are β-1,4-linked sugars, but xylan has a backbone of xylose rather than glucose units, and unlike cellulose is decorated with varied side chain substitutions (Qaseem et al., 2025). Hemicelluloses are synthesised in the Golgi apparatus and secreted into the wall during SCW deposition. As cells transition from primary to SCW formation, Golgi polysaccharide synthesis shifts from mainly pectin and xyloglucan to xylan and mannan (Meents et al., 2018). The synthesis, targeted secretion and assembly of xylan, in coordination with cellulose and lignin, together produce the highly organized architecture of SCWs. Increasing evidence suggests that the structure and organization of SCW polymers themselves may influence their patterned deposition (Takenaka et al., 2018; Pfaff et al., 2024). For example, the structure of xylan polymers has been shown to affect SCW patterns in tracheary elements induced to transdifferentiate from protoplasts isolated from SCW synthesis mutants (Pfaff et al., 2024). Yet, whether the structure and interactions of different SCW polymers contribute to SCW patterning – and how this is orchestrated in the native developmental context of distinct SCW-forming cell types – remains an open question.

Here we show that *C. hirsuta CESA7* is essential for cellulose synthesis in the end*b* SCWs of explosive fruit and is required for assembling wall polymers into highly ordered SCW patterns. In *cesa7* mutants, lignin and xylan deposition inititates normally but end*b* SCWs fail to adopt the correct pattern and layered organization during subsequent wall thickening. Cortical microtubules direct the formation of SCW-depleted domains along cell edges, generating the hinged SCW pattern of end*b* cells in explosive fruit. Disrupting microtubules abolishes this pattern and prevents fruit from exploding. Therefore, both microtubules and *CESA7* are required in *C. hirsuta* fruit to produce the specialized end*b* SCW pattern that drives the mechanics of explosive seed dispersal.

## Results

### Coordinated cellulose and lignin deposition in end*b* SCWs

To characterise the end*b* cell wall and its patterning in *C. hirsuta*, we used immunofluorescence labelling of fruit cross-sections with antibodies that target various cell wall polysaccharides. Xyloglucan and pectins with a high or low degree of methyl esterification are recognized by LM25, LM20 and LM19 antibodies, respectively (Verhertbruggen et al., 2009; Pedersen et al., 2012) and found only in the primary walls of end*b* cells (Fig. 1A). LM20 signal was depleted in regions of the lateral walls adjacent to the thick SCW (Fig. 1A), suggesting that primary wall modifications, such as partial pectin demethylesterification, may be associated with SCW formation in end*b* cells. Xylan is recognized by the LM11 antibody (McCartney et al., 2005) and labelled the entire end*b* SCW, consistent with xylans being the predominant hemicellulose in dicot SCWs (Scheller and Ulvskov, 2010) (Fig. 1A, Fig. S5A).

**Figure 1.**
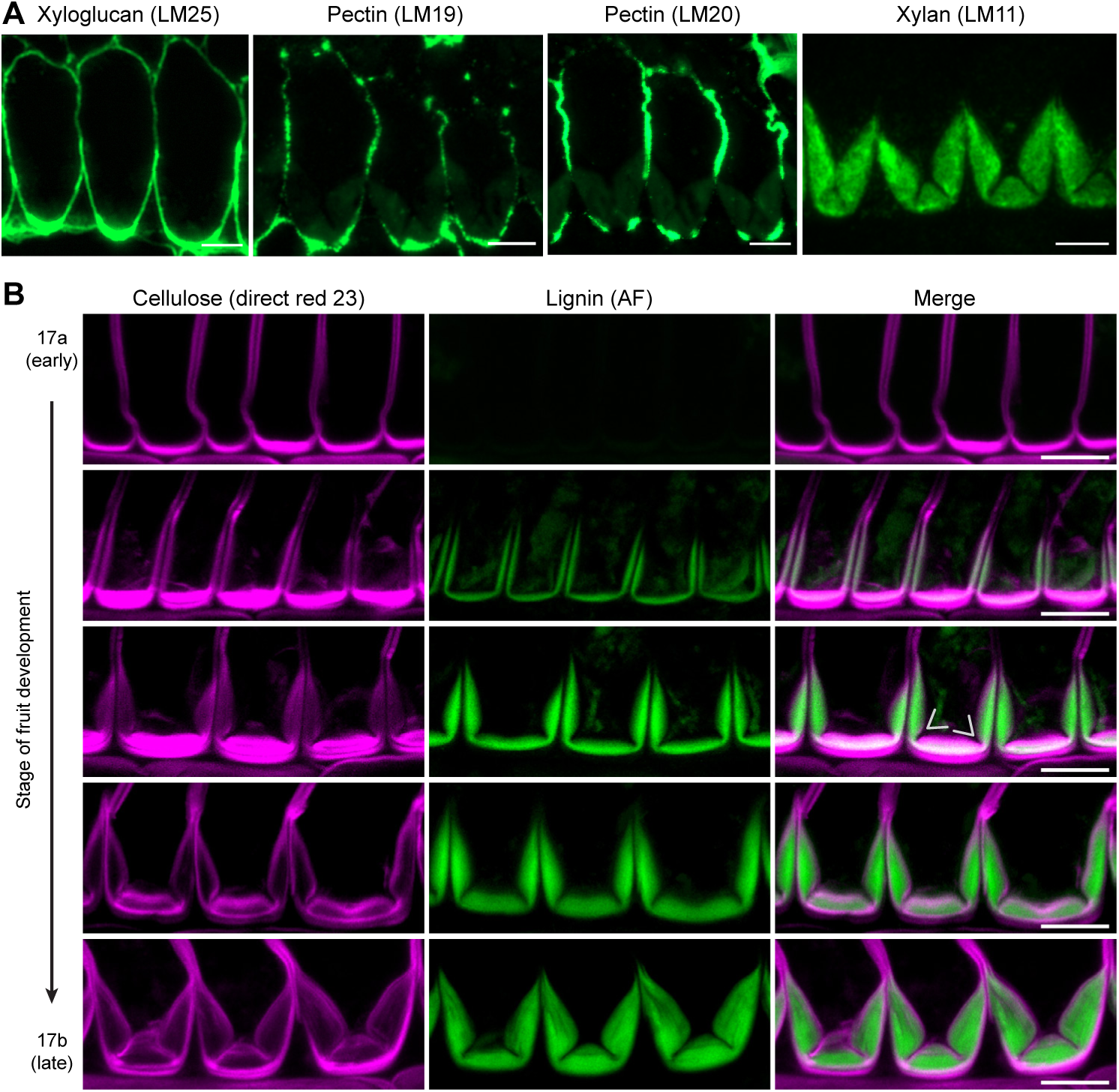
Coordinated cellulose and lignin deposition in *C. hirsuta* end*b* SCWs. **(A)** Immunofluorescence detection of cell wall epitopes in end*b* cells in transverse sections of resin-embedded *C. hirsuta* fruit at stage 17b. LM25 recognizes xyloglucans in the primary cell wall, LM19 and LM20 detect pectins with a low (LM19) or high (LM20) degree of methyl esterification in the primary cell wall, and LM11 recognizes xylan in the SCW. **(B)** Confocal laser scanning micrographs of end*b* SCW deposition during *C. hirsuta* fruit development from stages 17a through 17b showing cellulose stained with direct red 23 (magenta), lignin autofluorescence (AF, green) and both channels merged. Arrows indicate suppression of cellulose and lignin deposition along the cell edges to form two thin hinges that disrupt the thick SCW. Scale bars: 10 μm (A-B).

Cellulose thickening on the adaxial side of end*b* cells is an early indication of SCW formation (Fig. 1B). Given the abundance of xyloglucan and pectins in this region (Fig. 1A), it is possible that this thickening may include primary cell wall cellulose. This thickening slightly precedes a thin layer of lignin that is deposited in a “U”-shape on the adaxial side of end*b* cells (Fig. 1B). Cellulose and lignin deposition is immediately suppressed along the cell edges at the corners of this “U”, but continues throughout the rest of the SCW (Fig. 1B). This results in the formation of two thin hinges along the cell edges that disrupt the thick SCW (arrows, Fig. 1B). The coordinated deposition of cellulose and lignin throughout stage 17 of fruit development progressively thickens the SCW and enhances the hinge domain (Fig. 1B). Therefore, cellulose and lignin biosynthesis occur concurrently during patterning and formation of the hinged end*b* SCW. Given that the correct pattern of cellulose forms independently of lignin in the end*b* SCW of *lac4,11,17* mutants (Perez-Anton et al., 2022), this raises the question of whether patterned lignin deposition depends on cellulose.

### CESA7 controls cellulose biosynthesis in end*b* SCWs

To address this question, we took a genetic approach to perturb cellulose biosynthesis. We first screened Arabidopsis *CESA* mutant alleles for defects in end*b* SCWs. We observed almost no SCW cellulose and a slight reduction in lignin in the end*b* SCWs of the *CESA7* allele *irx3-4* (Brown et al., 2005) (Fig. 2A). By contrast, the end*b* SCW appeared wild type in the *CESA3* allele *cev1* (Ellis et al., 2002) (Fig. S1A-B). Therefore, *CESA7* is required for cellulose biosynthesis in the end*b* SCW in Arabidopsis. To test whether this function of *CESA7* is likely to be conserved in *C. hirsuta*, we complemented *irx3-4* with the *C. hirsuta CESA7* cDNA expressed under its own promoter, tagged at the N-terminus with mNeonGreen (*pCESA7::mNG:CESA7*). Cellulose biosynthesis was fully restored in the end*b* SCW of complemented *irx3-4* fruit, indicating a high degree of conservation in *CESA7* function between *C. hirsuta* and Arabidopsis (Fig. 2A). To analyze *CESA7* expression in *C. hirsuta*, we generated a transcriptional reporter (*pCESA7::3xGFP:NLS*) and found expression in end*b* cells and other cells with SCWs, such as protoxylem cells in the root (Fig. 2B, Fig. S3B). We found that *C. hirsuta CESA7* localized to end*b* cells before and during SCW formation (Fig. 2B), suggesting that *CESA7* may be required for cellulose biosynthesis in the end*b* SCW of both Arabidopsis and *C. hirsuta* fruit.

**Figure 2.**
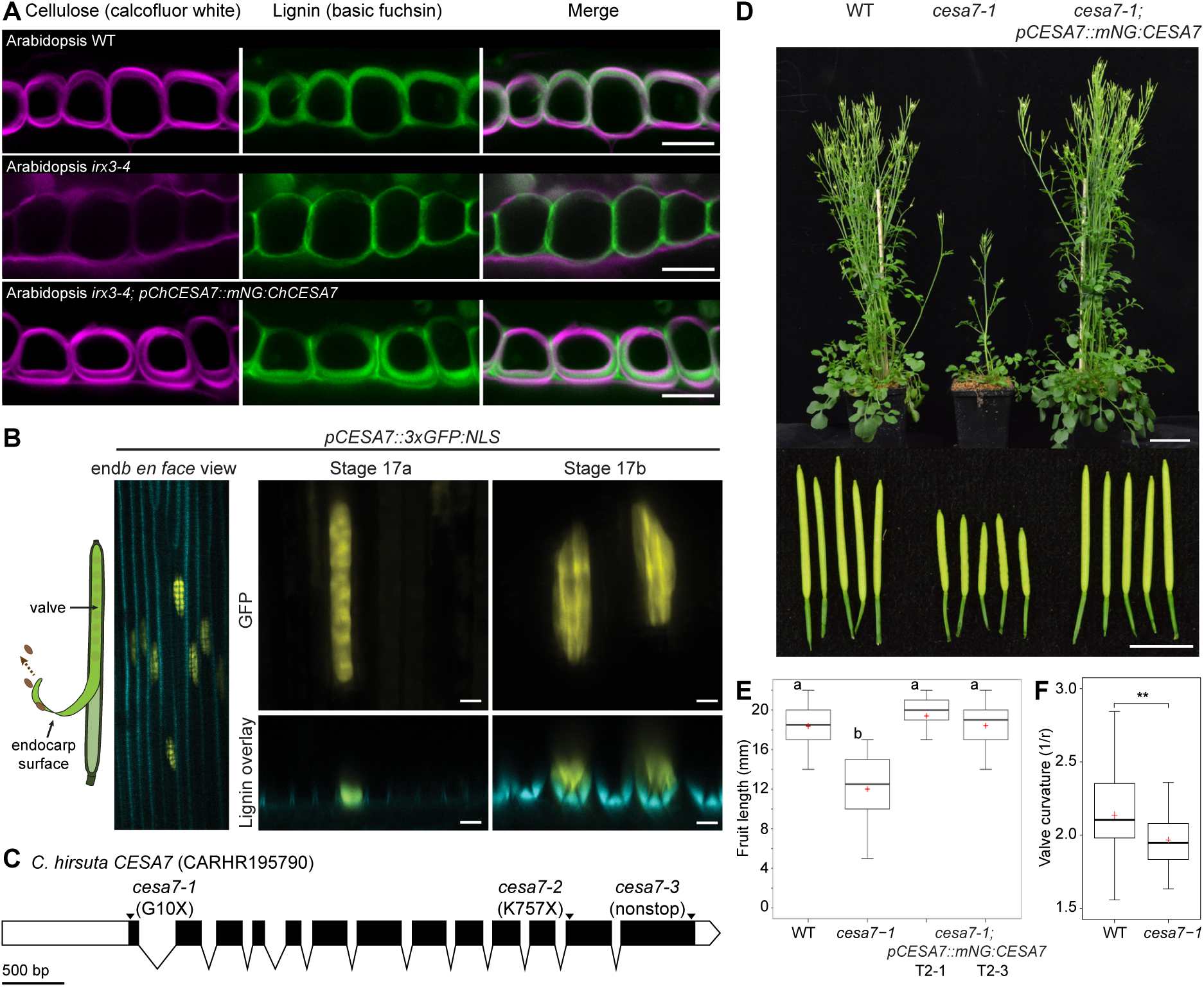
*cesa7* mutant alleles in *C. hirsuta*. **(A)** Confocal laser scanning micrographs of end*b* SCWs in Arabidopsis stage 17b fruit showing cellulose stained with calcofluor white (magenta), lignin stained with basic fuchsin (green) and both channels merged in wild type, *irx3-4* (*CESA7* allele) and *irx3-4* complemented with a *C. hirsuta pCESA7::mNG:CESA7* fusion protein. **(B)** Expression of *C. hirsuta pCESA7::3*×*GFP:NLS* (yellow) in *C. hirsuta* end*b* cell nuclei of stage 17a and 17b fruit valves imaged *en face* (left and upper panels), transverse optical sections (lower panels) shown overlaid with lignin autofluorescence (cyan). Schematic indicates the endocarp surface of the fruit valve used for imaging. **(C)** Gene model of *C. hirsuta CESA7* showing the *cesa7-1*, *cesa7-2* and *cesa7-3* CRISPR/Cas9 mutations together with the location of sgRNAs (black triangles), UTRs (white bars), exons (black bars) and introns (lines). Scale bar: 500 bp. **(D)** Whole plant and fruits of *C. hirsuta* wild type, *cesa7-1* and *cesa7-1* complemented with a *C. hirsuta pCESA7::mNG:CESA7* fusion protein. **(E)** Boxplot of fruit length in wild type, *cesa7-1* and two independent *cesa7-1; pCESA7::mNG:CESA7* complemented lines (n = 50 fruit from 5 different plants per genotype). Different letters denote statistical significance at *p* = 6.047e-30 using ANOVA followed by Tukey’s Honestly Significant difference (HSD) test. **(F)** Boxplot of coiled valve curvature (1/radius) in wild-type (n = 33) and *cesa7-1* (n = 39) fruit. ** denotes statistical significance at *p* = 0.01 using Wilcoxon rank sum test. All plots show median (thick black line) and mean (red cross). Scale bars: 10 μm (A-B), 5 cm (plants, D), 5 mm (fruit, D).

To perturb *CESA7* function in *C. hirsuta*, we generated *CESA7* mutant alleles by CRISPR/Cas9 gene editing. We recovered three recessive alleles, including *cesa7-1* where an 8-bp deletion resulted in a truncated ten-amino acid protein lacking all functional domains (Fig. 2C, Fig. S2). Mature *cesa7-1* plants were dwarfed with significantly shorter fruits than wild-type, similar to Arabidopsis *irx3-4* (Fig. 2D-E, Fig. S1C-D). We assessed the explosive coiling of *cesa7-1* fruit valves by measuring the curvature of exploded valves, since more tightly coiled valves reflect more explosive energy release (Hofhuis et al., 2016). We found a significant reduction in *cesa7-1* valve curvature compared to wild type (Fig. 2F), indicating that valve coiling is less explosive. Phenotypes of *cesa7-1* were fully complemented by expressing wild-type *C. hirsuta CESA7* (*pCESA7::mNG:CESA7*) (Fig. 2D-E, Fig. S3A). Therefore, *cesa7-1* represents loss of *CESA7* function in *C. hirsuta*.

We could detect very little cellulose in the end*b* SCW of *cesa7-1* fruit using direct red 23 as a cellulose-specific dye (Fig. 3A, Fig. S3C). In contrast, cellulose was clearly stained in the primary walls of *cesa7-1* end*b* cells (Fig. 3A). We quantified crystalline cellulose in *C. hirsuta* fruit valves and found that *cesa7-1* mutants retained only 46.9% of the cellulose present in wild-type valves (Fig. 3B). This suggests that more than half of the cellulose content of *C. hirsuta* fruit valves is found in SCWs; mostly contributed by the end*b* cell layer. Reduced cellulose also resulted in the collapse of xylem vessels in the replum of *cesa7-1* fruits (Fig. S3D), similar to Arabidopsis *cesa7* mutants (Taylor et al., 1999; Brown et al., 2005; Takenaka et al., 2018). Therefore, *CESA7* is required for cellulose biosynthesis in SCWs of *C. hirsuta* fruit.

**Figure 3.**
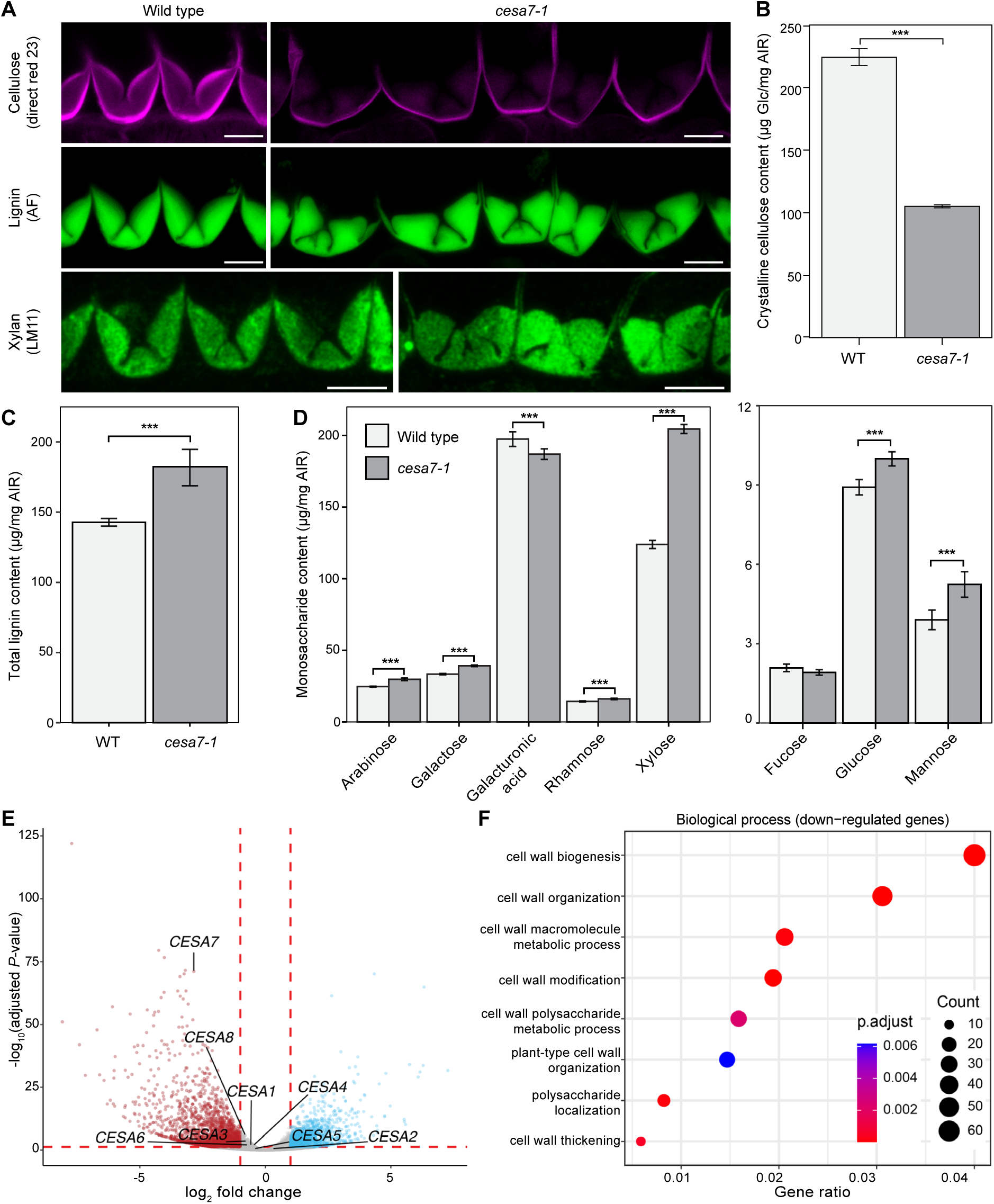
CESA7 controls cellulose biosynthesis in *C. hirsuta* end*b* SCWs. **(A)** *C. hirsuta* wild-type and *cesa7-1* end*b* SCWs in stage 17b fruit showing cellulose stained with direct red 23 (magenta), lignin autofluorescence (green) and immunofluorescence detection of xylan using LM11 antibody. **(B-D)** Boxplots of crystalline cellulose, lignin and matrix monosaccharides in mature fruit valves of wild type vs *cesa7-1*, shown as μg per mg of alcohol-insoluble residue (AIR) for glucose (B), acetylbromide-soluble lignin (C), and matrix polymer sugars (D). Bars indicate mean values; error bars represent ± standard deviation calculated from 4 biological replicates of pooled valves (approximately 530 valves were pooled for wild type and 660 valves for *cesa7-1*); *** denotes statistical significance at *P* < 0.001 using Student’s t-test. **(E)** Volcano plot of differential gene expression between wild-type and *cesa7-1* fruit valves (logFC ≥ |1|, FDR < 0.05) showing 1327 up-regulated genes (blue) and 2138 down-regulated genes (red). All *CESA* genes expressed in *C. hirsuta* fruit valves are indicated; only *CESA7* is differentially expressed. **(F)** Selected gene ontology terms enriched in the 2138 down-regulated genes. Scale bars: 10 μm (A).

We also treated *C. hirsuta* fruit with the cellulose synthesis inhibitors DCB (2,6-dichlorobenzonitrile) and isoxaben. However, neither chemical affected end*b* SCW biosynthesis in our experimental conditions (Fig. S4A). Isoxaben treatment affected only primary cell walls in the fruit (Fig. S4B-C), consistent with its specificity towards cellulose synthase complexes in primary, rather than SCWs (Larson and McFarlane, 2021). Therefore, by taking a genetic approach, we were able to substantially reduce SCW cellulose and identify a critical role for *CESA7* in cellulose synthesis in the end*b* SCW of both Arabidopsis and *C. hirsuta* fruit.

### Patterned lignin and xylan deposition is independent of cellulose

Despite the depletion of cellulose, end*b* SCWs are fully lignified in *cesa7* fruit (Fig. 3A). We used LM11 antibody labelling to show that the hemicellulose xylan is deposited in the same hinged pattern as lignin in *cesa7* end*b* SCWs (Fig. 3A). Therefore, the spatial deposition of lignin and xylan is not strictly dependent on cellulose patterning in end*b* SCWs of *C. hirsuta* fruit. The abundance of hemicellulose monosaccharides and lignin were significantly increased in *cesa7-1* compared to wild-type fruit valves (Fig. 3C-D). In particular, the proportion of xylose, indicative of xylan, was significantly increased in *cesa7-1* (Fig. 3D). We observed elevated glucose in *cesa7-1* despite the significant reduction in crystalline cellulose, which may indicate an increase in non-crystalline, amorphous cellulose and/or primary wall xyloglucans (Fig. 3D). Primary cell wall pectic polysaccharides, represented by galacturonic acid, were reduced in *cesa7-1* (Fig. 3D). Overall, these findings indicate that *cesa7* fruit valves contain significantly less crystalline cellulose and more lignin and hemicellulose than wild-type, and that cellulose synthesis does not pre-pattern the hinged geometry of end*b* SCWs.

To investigate how these differences in cell wall composition were reflected at the transcriptome level, we compared RNAseq data from *cesa7-1* versus wild-type fruit valves. We identified 3465 differentially expressed genes based on a false discovery rate (FDR) of <0.05 and a minimum fold-change of 2, with 1327 up-regulated and 2138 down-regulated genes in *cesa7-1* (Table S1). *CESA7* is among the top five most significantly down-regulated genes in *cesa7-1* valves (Fig. 3E, Table S1). The other seven *CESA* genes expressed in valves were not differentially expressed in *cesa7-1*, suggesting no co-regulation or compensatory regulation of these genes in response to loss of *CESA7* (Fig. 3E). Although xylan levels were proportionally higher in *cesa7-1* compared to wild-type cell walls (Fig. 3D), we did not observe upregulation of xylan synthesis genes (Table S1), suggesting that post-transcriptional processes may contribute to this increase. Gene Ontology (GO) analysis showed that down-regulated genes in *cesa7-1* fruit valves were enriched for cell wall biological processes (Fig. 3F, Table S1), suggesting that cell wall biogenesis and organization is generally perturbed in the absence of *CESA7*.

### Failure to maintain SCW patterning in *cesa7* mutants

In *cesa7* mutants, the end*b* SCW retains key features of the wild-type pattern: deposition remains polar, and hinges form along the cell edges, indicating that xylan and lignin are initially deposited in the correct domain (Fig. 3A). Nevertheless, the mature end*b* SCW pattern deviates from the wild type in all three *cesa7* alleles (Fig. 3A, Fig. S3E). To determine whether these defects arise from a failure to correctly initiate or to maintain the SCW pattern, we analyzed a developmental time course of end*b* SCW formation in *cesa7-1* fruit (Fig. 4A). We could only detect cellulose in the primary wall of *cesa7* end*b* cells, and observed the same initial thickening on the adaxial side of cells that we observed in wild type (Fig. 1B, Fig. 4A). Lignin is deposited in a thin, “U”-shaped pattern on this adaxial side, followed by the initiation of hinges along the cell edges at the corners of this “U” (arrows, Fig. 4A). Therefore, the distinctive pattern of the end*b* SCW appears to initiate normally, despite the absence of detectable cellulose in the SCW. The hinge domains are enhanced as the rest of the lignified wall thickens (Fig. 4A). During this thickening, the hinged regions become wider and the SCW surface undulates (Fig. 1B, Fig. 4A). The mature SCW is misshapen compared to wild type with the precise pattern varying from cell to cell (Fig. 4A). Therefore, *cesa7* end*b* cells fail to maintain SCW patterning during subsequent thickening.

**Figure 4.**
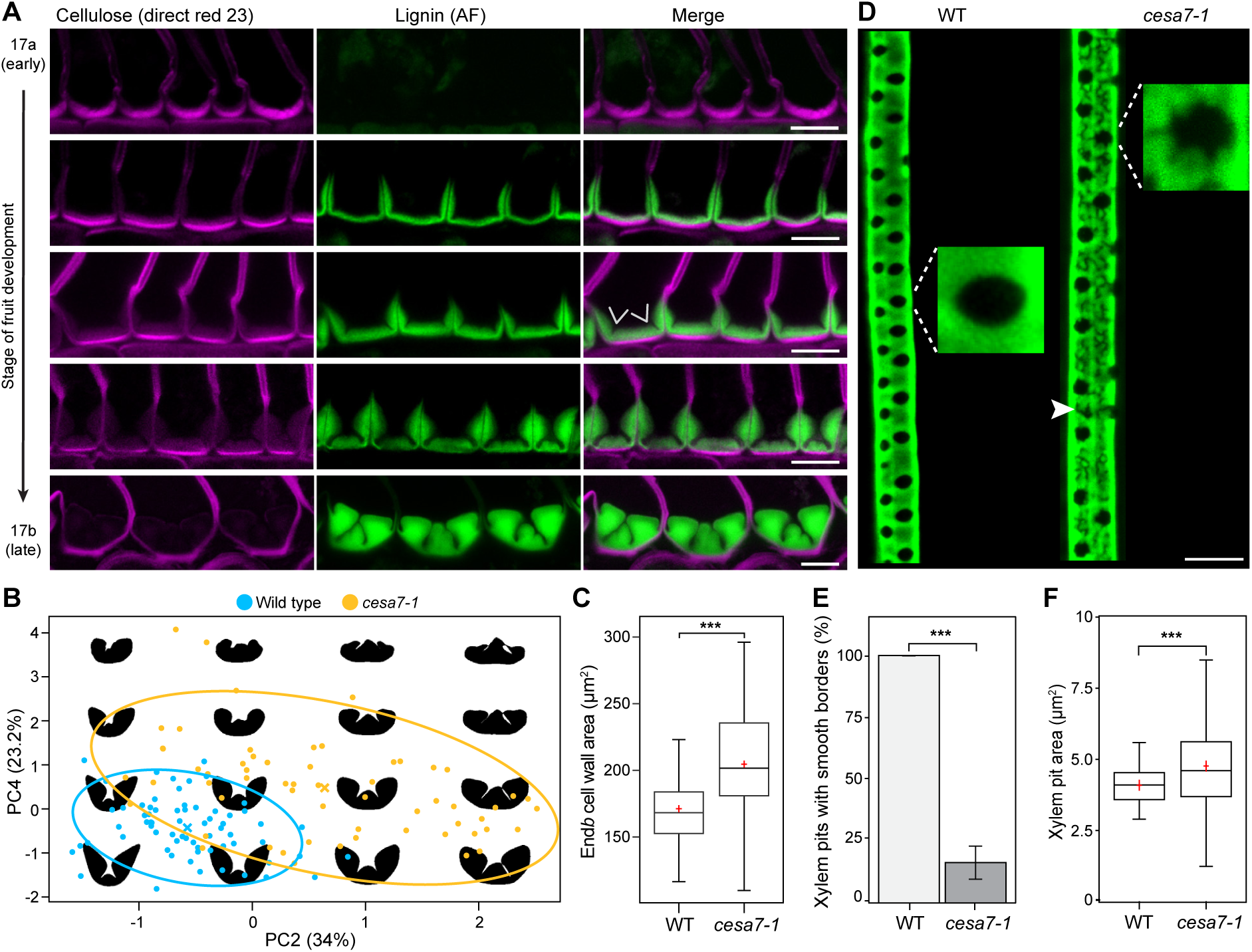
CESA7 is required to maintain SCW patterning in end*b* and metaxylem cells in *C. hirsuta*. **(A)** Confocal laser scanning micrographs (CLSM) of end*b* SCW deposition in *cesa7-1* fruit during stages 17a through 17b showing cellulose stained with direct red 23 (magenta), lignin autofluorescence (AF, green) and both channels merged. Arrows indicate suppression of cellulose and lignin deposition along the cell edges to form a hinged SCW pattern. **(B)** Shape-space plot of end*b* SCW contours of wild type and *cesa7-1*, based on principal component analysis (PCA). PC2 and PC4 values for wild-type (blue) and *cesa7-1* (yellow) contours are plotted along the x and y axes as multiples of their respective standard deviations (% of explained variance is indicated). Crosses indicate genotype means; ellipses indicate half the standard deviation. **(C)** Boxplot of end*b* SCW cross-sectional area in wild type and *cesa7-1* fruit (n = 75 end*b* cells in WT and 83 end*b* cells in *cesa7-1*). **(D)** Maximum projection of CLSM stacks of metaxylem SCWs in wild type and *cesa7-1* roots in 6-day old seedlings showing lignin stained with basic fuchsin (green). Zoom-in of a single pit is shown for each genotype. **(E)** Barplot showing the percentage of metaxylem pits with smooth borders in wild-type and *cesa7-1* roots. Bars indicate mean values; error bars represent ± standard deviation (n = 280 pits in 9 WT roots and 359 pits in 13 *cesa7-1* roots). **(F)** Boxplot of metaxylem pit cross-sectional area in wild-type and *cesa7-1* roots (n = 89 pits in WT and 319 pits in *cesa7-1*). *** denotes statistical significance at *P* < 0.001 using Student’s t-test (C, E-F). Boxplots show median (thick black line) and mean (red cross). Scale bars: 10 μm (A, D).

To quantify the variation in end*b* SCW patterning in *cesa7* versus wild-type fruit, we used the multivariate shape analysis tool LeafI (Zhang et al., 2020). We used this software as previously described to extract contours of end*b* SCWs from fruit cross sections, specify landmarks, and perform shape space analysis and visualization (Zhang et al., 2020). Using principal-component analysis (PCA), we found two PC axes that accounted for 57.2% of the variance and discriminated the broad distribution of SCW shapes found in *cesa7* from the restricted shape space of wild-type SCWs (Fig. 4B). Shape models along these axes captured the irregular overgrowth of the *cesa7* SCW, which was accompanied by a significant increase in end*b* SCW area compared to wild type (Fig. 4B-C). Thus, although the initial pattern of xylan and lignin deposition is preserved in *cesa7* end*b* cells, the assembly of these polymers in the cellulose-deficient SCW appears disrupted, resulting in aberrant SCW patterns.

To test whether *CESA7* is required more generally to maintain SCW patterning in other cell types, we examined metaxylem cells in the roots of 6-day old *C. hirsuta* wild-type and *cesa7* seedlings. The patterning of SCW pits in metaxylem cells is a well-characterized process in Arabidopsis (Oda and Fukuda, 2012b, a; Xu et al., 2022). In wild-type *C. hirsuta* roots, metaxylem cells had circular/oval pits with smooth borders (Fig. 4D). In *cesa7* metaxylem SCWs, pits were significantly larger on average and displayed high variability compared to wild type (Fig. 4F). Most strikingly, the vast majority of pits in *cesa7* had irregular borders, compared to the smooth borders of wild-type pits (Fig. 4D-E). The lignified SCW was also patchy in *cesa7* metaxylem cells, indicating that the continuous distribution of lignin within the cell wall matrix relies on SCW cellulose (arrowhead, Fig. 4D). Therefore, *CESA7* is necessary to maintain the pattern of SCW pits in metaxylem cells and end*b* SCWs in *C. hirsuta* fruit.

### Highly ordered pattern and layered architecture of end*b* SCWs depends on *CESA7*

Since two-dimensional fruit cross sections do not fully capture the shape of the end*b* SCW, we analyzed wild-type and *cesa7* end*b* SCWs in three dimensions. We acquired z-stacks of the entire lignified SCW of long end*b* cells in intact fruit valves and used these to generate three-dimensional renderings with Imaris software. The surface of the wild-type SCW was continuous and smooth along the cell length (Fig. 5A). In contrast to wild type, the surface of the *cesa7* SCW was undulated with outgrowths and indentations along the entire cell length (Fig. 5B). This explains why the end*b* SCW pattern varied between different cross-sections of the same cell, and between different cells in *cesa7* fruit (Fig. 4A-B). Maximum-intensity projections of the confocal z-stacks showed that these undulations created a patchy end*b* SCW with an uneven distribution of lignin in *cesa7* fruit (Fig. 5C-D). Therefore, cellulose is required to produce a regular, highly ordered SCW pattern in end*b* cells.

**Figure 5.**
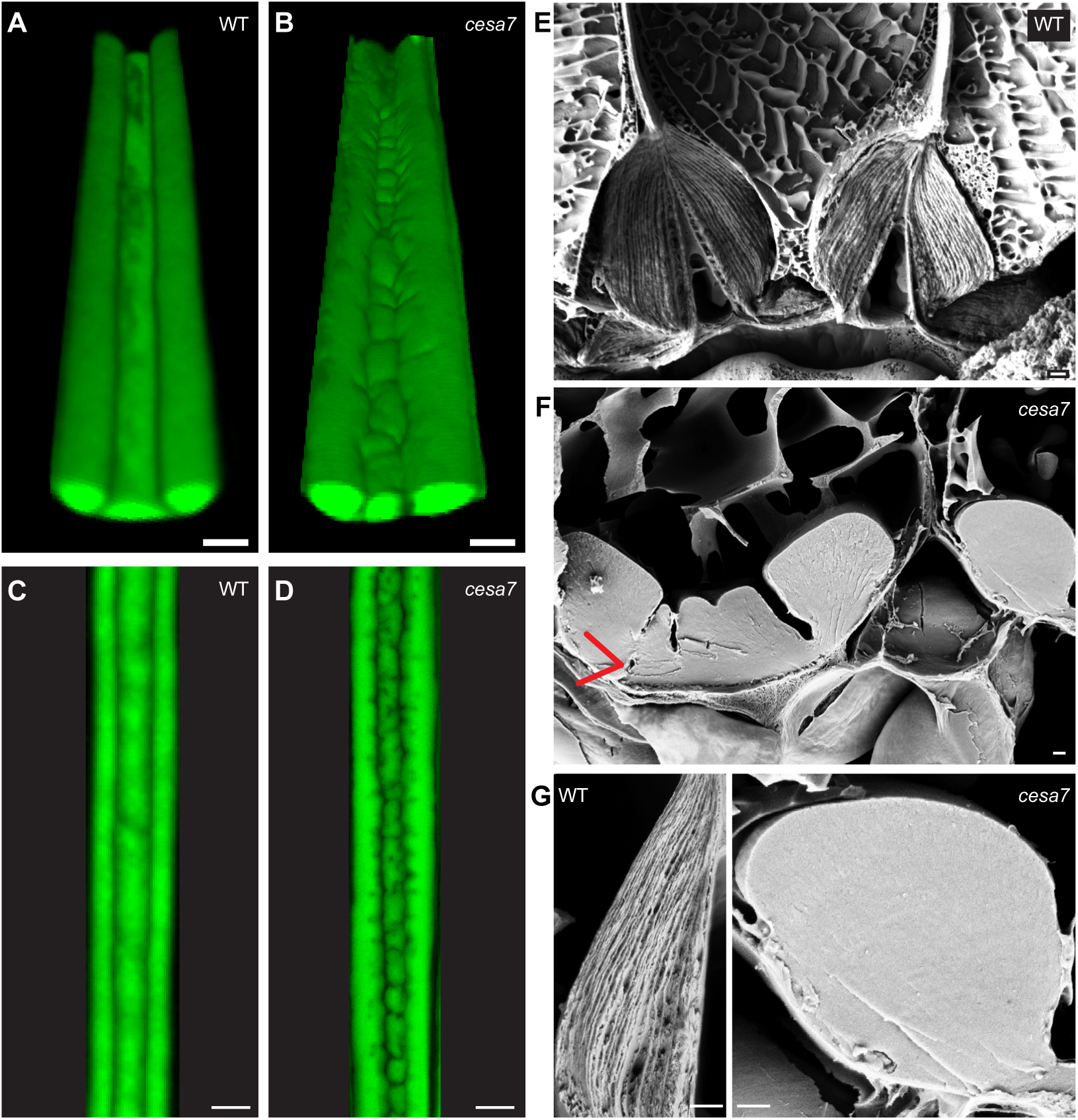
The regular fibrous arrangement of lignocellulosic material in wild-type end*b* SCWs is replaced by dense undulating cell wall material in *C. hirsuta cesa7-*1. **(A-D)** Two-photon excitation laser scanning microscopy of intact *C. hirsuta* fruit valves showing lignin autofluorescence (green) of end*b* SCWs in wild type and *cesa7-1* as three-dimensional renderings using Imaris software (A-B) and maximum projections (C-D). **(E-G)** Cryo-fracture scanning electron micrographs of *C. hirsuta* wild type and *cesa7-1* end*b* SCWs. Red arrow indicates hole in SCW where initiated hinge was subsequently filled (F). Scale bars: 10 μm (A-D), 1 μm (E-G).

To compare the architecture of wild-type end*b* SCWs with the cellulose-deficient SCWs in *cesa7* end*b* cells, we used cryo-fracture scanning electron microscopy. The lignocellulosic material in wild-type end*b* SCWs is fibrous and organized into many fine layers that form the regular, hinged SCW shape (Fig. 5E, G). In stark contrast, the *cesa7* end*b* SCW material is non-fibrous and lacks this layered organization (Fig. 5F-G). The condensed material in *cesa7* end*b* SCWs instead forms irregular, hinged shapes with undulating folds. We observed holes within the SCW where a hinge appeared to initiate, but was subsequently overgrown by newly deposited SCW material (arrow, Fig. 5F). Therefore, SCW polymers in *cesa7* end*b* SCWs fail to assemble into a layered structure and cannot maintain a highly ordered SCW pattern.

### Microtubules direct end*b* SCW patterning

Although the end*b* SCW is aberrant in *cesa7*, hinges remain patterned along the cell edges, allowing the fruit valves to coil, albeit less explosively than in wild type (Fig. 2F, 4A). To investigate the mechanisms underlying this patterning, we examined the role of cortical microtubules. We used two-photon excitation microscopy to visualize the microtubule marker GFP-TUA6 (Mosca et al., 2024) in wild-type end*b* cells during SCW development. Despite the large size of end*b* cells and their deep tissue location within the fruit valve, we were able to image microtubules associated with the SCW on the adaxial and adjacent lateral sides of each cell. We used Imaris software to visualize microtubules in optical slices through both the adaxial and adjacent lateral side of end*b* cells in valves of stage 16 and 17a fruit (Fig. 6A-B). Cortical microtubule arrays were dense and well-organized at these stages, aligning transversely to the long axis of the cell (Fig. 6A-B), consistent with conventional mechanisms for anisotropic growth in these extremely long cells. Following the initial deposition of the SCW at stage 17a (Fig. 6B), fruit elongation ceased, and by stage 17b, the microtubule arrays had adopted a more longitudinal orientation (Fig. 6C). As SCW thickening progressed during stage 17b, the cortical space became restricted, and microtubules were positioned along the cell cortex adjacent to the SCW thickenings (Fig. 6C-D). However, these images lacked the resolution necessary to discriminate any patterning information about the distribution of microtubules along cell edges where local SCW depletion forms the hinged SCW pattern.

**Figure 6.**
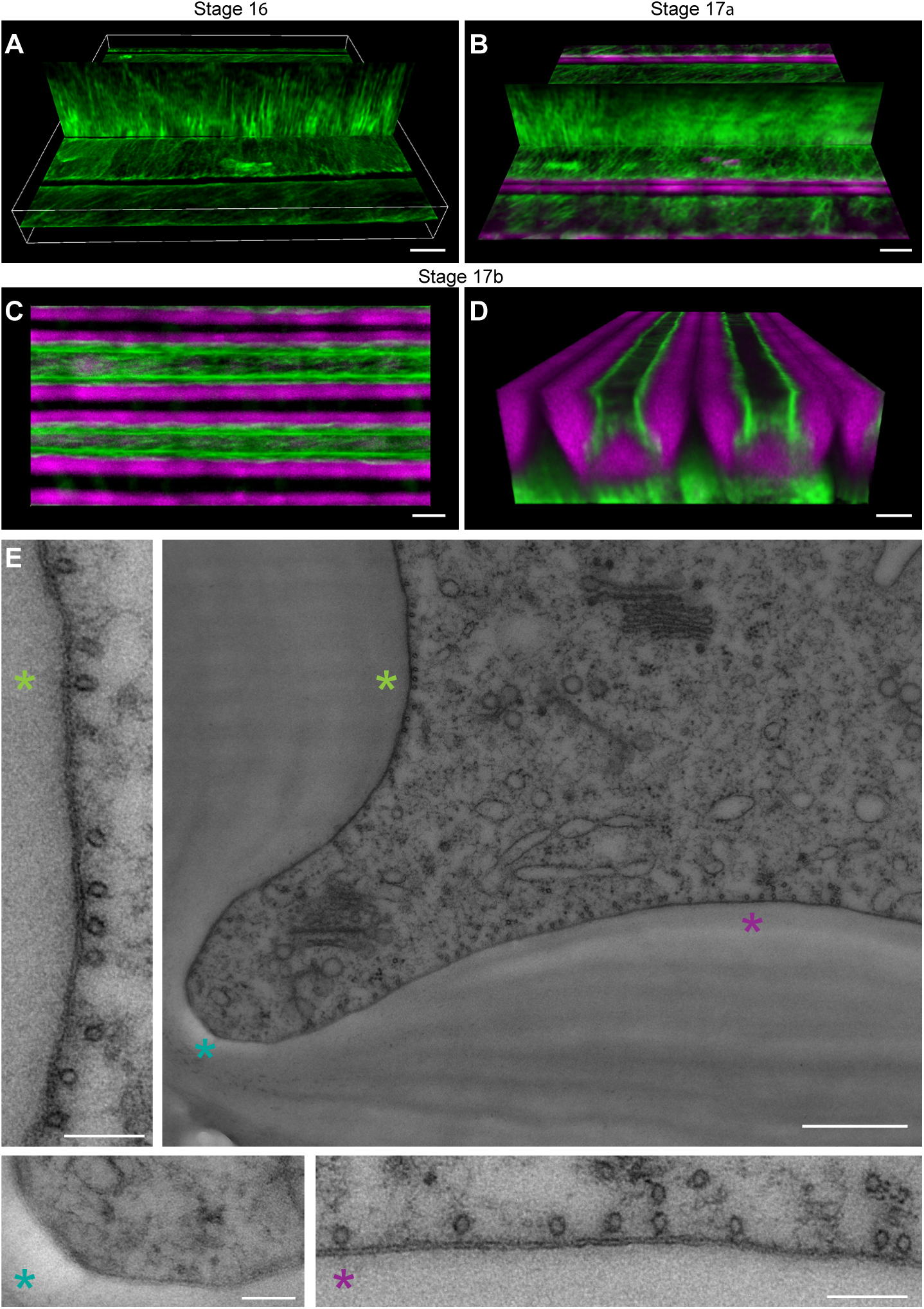
Cortical microtubule density is high at regions adjacent to SCW thickenings and low at regions of SCW depletion in *C. hirsuta* end*b* cells. **(A-D)** Two-photon excitation laser scanning microscopy of intact *C. hirsuta 35S::GFP:TUA6* fruit valves showing GFP:TUA6 signal (green) and lignin autofluorescence (magenta) in end*b* cells at stages 16 (A), 17a (B) and 17b (C-D). **(E)** Transmission electron micrograph of a cross section of a high-pressure frozen *C. hirsuta* fruit showing the hinge domain of an end*b* SCW. A zoom-in of each of the three regions indicated by a coloured asterisk (*) is shown. Scale bars: 5 μm (A-D), 0.5 μm (large panel, E), 0.1 μm (small panels, E).

To obtain high resolution sub-cellular views of cortical microtubule distribution adjacent to the end*b* SCW, we used transmission electron microscopy of cryo-immobilized fruit cross-sections. We observed a high density of microtubules lining the plasma membrane in regions adjacent to SCW thickenings and a low microtubule density at the cell edges, adjacent to areas where SCW depletion forms a hinge (Fig. 6E). This distribution is suggestive of a patterning role for cortical microtubules whereby the local density of microtubules in the adjacent cortex may direct SCW deposition.

To test whether cortical microtubules are required to pattern the hinged end*b* SCW, we disrupted microtubules using two complementary approaches: treatment with the microtubule depolymerizing drug oryzalin and inducible expression of a truncated version of the atypical tubulin kinase PROPYZAMIDE-HYPERSENSITIVE 1 (PHS1ΔP) (Fujita et al., 2013) (*pUBQ10::GR-LhG4/pOp6::PHS1ΔP:mCherry*). Dexamethasone (Dex) induction of *PHS1ΔP:mCherry* expression resulted in full depolymerization of microtubules in end*b* cells (Fig. 7A). Analysis of cross-sections revealed that both oryzalin-treated fruit and Dex-induced *pUBQ10::LhGR>>PHS1ΔP:mCherry* fruit completely lost the hinged pattern in end*b* SCWs (Fig. 7B, Fig. S6A-B). While mock-treated fruit showed a normal hinged end*b* SCW, both oryzalin-treated and Dex-induced fruits had thickened SCWs that lacked the hinge domains (Fig. 7B, Fig. S6A-B). This loss of patterning was apparent in the deposition of cellulose, lignin, and xylan in the end*b* SCWs of Dex-induced *pUBQ10::LhGR>>PHS1ΔP:mCherry* fruit (Fig. 7B-C, Fig. S5B). Moreover, the lignified SCW extended around the entire end*b* cell in Dex-treated compared to mock-treated fruit (Fig. 7B). Therefore, cortical microtubules may direct the delivery of all SCW components to the plasma membrane in the precise pattern required to produce a polar, hinged SCW in *C. hirsuta* end*b* cells.

**Figure 7.**
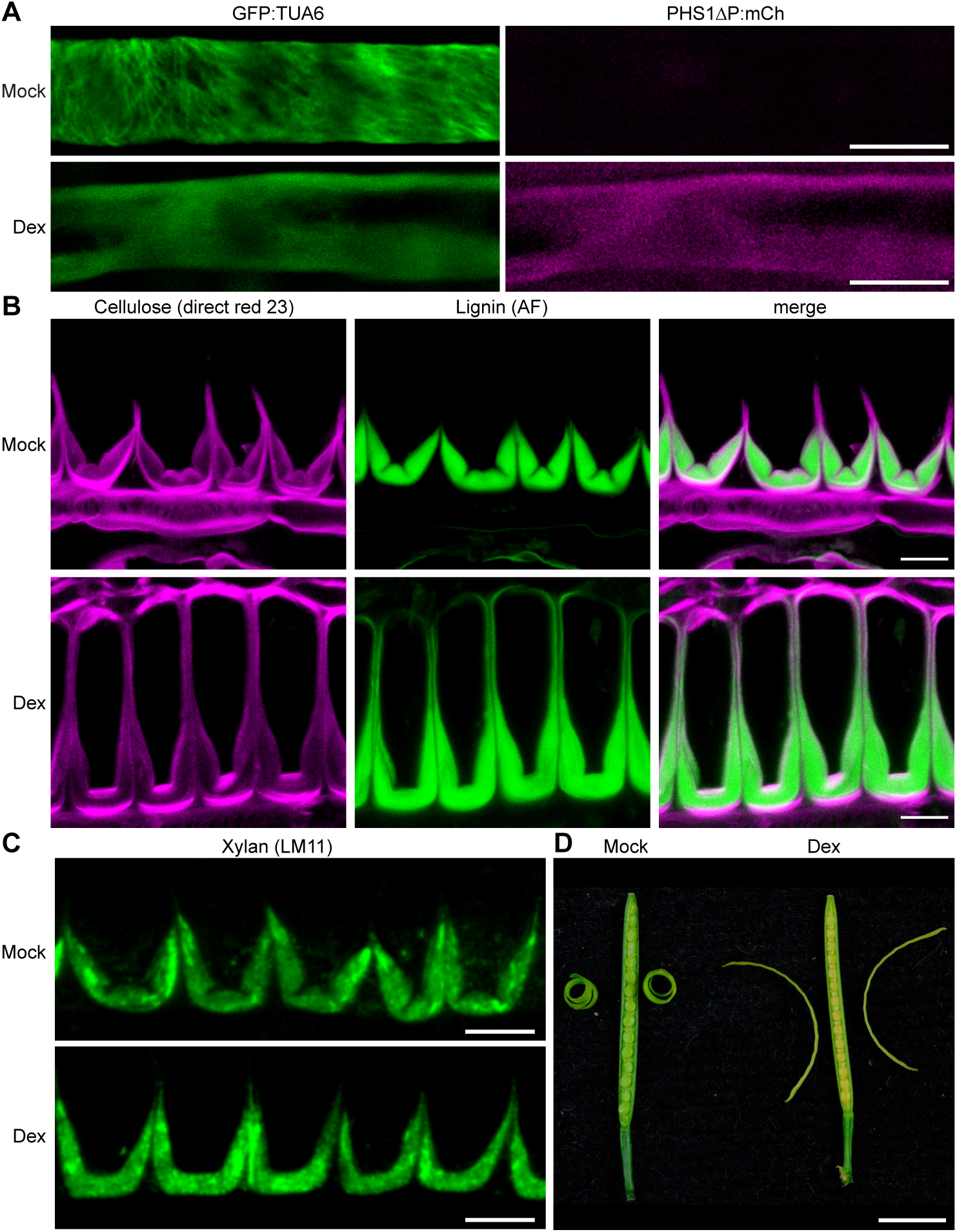
Microtubules are necessary for end*b* SCW patterning and explosive valve coiling in *C. hirsuta* fruit. **(A-D)** *C. hirsuta pUBQ10::GR-LhG4/pOp6::PHS1ΔP:mCherry; p35S::GFP:TUA6* fruit grown for 11 days on media without (mock) or with 1 mM dexamethasone (Dex). (A) Confocal laser scanning micrographs (CLSM) of end*b* cells in fruit after 5 days growth on media, imaged *en face* showing GFP-TUA6 signal (green) and PHS1ΔP-mCherry signal (magenta). (B) CLSM showing cellulose stained with direct red 23 (magenta), lignin autofluorescence (AF, green) and both channels merged in end*b* SCW cross sections of stage 17b fruit after 11 days growth on media. (C) Immunofluorescence detection of xylan using LM11 antibody (green) in end*b* SCW cross sections of resin-embedded stage 17b fruit after 11 days growth on media. (D) Exploded fruit after 10 days of daily treatment with mock or 2 mM Dex solution. Scale bars: 5 μm (A), 10 μm (B-C), 5 mm (D).

To assess the functional consequences of this loss of SCW patterning, we examined explosive valve coiling in mature fruit. Both oryzalin-treated fruit and Dex-induced *pUBQ10::LhGR>>PHS1ΔP:mCherry* fruit were non-explosive, and the valves failed to coil, whereas mock-treated fruit exploded normally with tightly coiled valves (Mosca et al., 2024) (Fig. 7D). These results indicate that cortical microtubules are required to pattern the polar, hinged SCW in *C. hirsuta* end*b* cells, which in turn is essential for explosive valve coiling.

### End*b* SCW patterning depends on both microtubules and CESA7

We next examined the effects of simultaneously losing both microtubules and *CESA7* on the hinged end*b* SCW pattern. To this end, we depolymerized microtubules in *cesa7* fruit using oryzalin (Fig. 8A). Cross sections of these fruits showed a complete loss of the hinged pattern in end*b* SCWs (Fig. 8A). Surprisingly, the aberrant end*b* SCW in *cesa7* was considerably enhanced by the loss of microtubules. In place of three distinct SCW domains separated by hinges along the cell edges, a very thick, amorphous SCW formed on the adaxial side of end*b* cells (Fig. 8A). Thus, overgrowth of the cellulose-depleted SCW appears to be amplified in the absence of microtubule patterning. By analyzing *cesa7* fruit where the oryzalin treatment was less consistent, we observed holes within the SCW where a hinge had been initiated, but not maintained, and subsequently overgrown by newly deposited SCW layers (Fig. S6C-E). In summary, the organized distribution of microtubules directs the formation of SCW-depleted domains along end*b* cell edges, establishing a hinged SCW pattern (Fig. 8B). Cellulose maintains this pattern by providing a layered scaffold that supports the compact assembly of SCW polymers (Fig. 8B). In the absence of both microtubule organization and normal crystalline cellulose deposition, SCWs are overgrown with no patterning (Fig. 8B). Therefore, both cortical microtubules and *CESA7* are required in *C. hirsuta* fruit to produce the mature end*b* SCW pattern.

**Figure 8.**
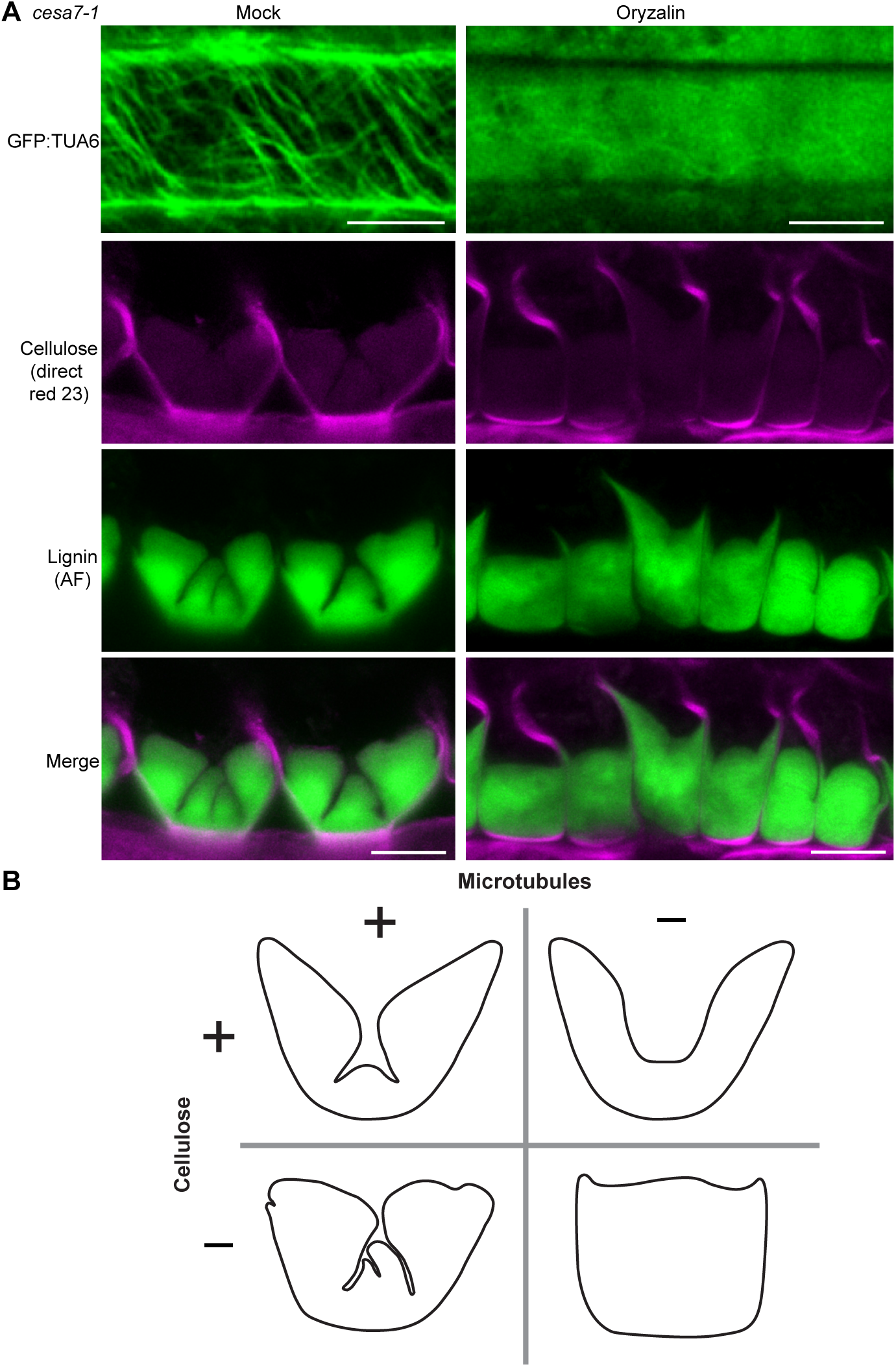
Microtubule depolymerization amplifies the overgrowth of cellulose-depleted end*b* SCWs in *C. hirsuta cesa7-1*. **(A)** *C. hirsuta cesa7-1; p35S::GFP:TUA6* stage 17b fruit after mock or oryzalin treatment, imaged *en face* showing GFP-TUA6 signal (green) or in cross section showing cellulose stained with direct red 23 (magenta), lignin autofluorescence (AF, green) and both channels merged. **(B)** Cartoon summarizing the effects of removing cellulose (*cesa7*) and/or depolymerizing cortical microtubules on the resulting pattern of end*b* SCWs in *C. hirsuta* fruit. Scale bars: 5 μm (A).

## Discussion

Explosive seed dispersal relies on the distinctive SCW pattern of end*b* cells in *C. hirsuta* fruit valves. We showed here that *CESA7* is required to synthesize the large amounts of cellulose in these specialized SCWs. Although SCW patterning can be initiated, it cannot be correctly maintained in *cesa7* end*b* cells, suggesting that cellulose scaffolds the proper assembly of SCW polymers into a layered organization that is essential for pattern formation. Cortical microtubules, in turn, are required to initiate the precise patterning of thin hinges that disrupt the thick end*b* SCW and enable explosive coiling of the fruit valves. In this way, both microtubules and *CESA7* contribute to the form and function of end*b* SCWs in exploding seed pods.

Mature end*b* cells in *C. hirsuta* have a typical primary wall composed of pectins, xyloglucan, and cellulose synthesized by primary wall CSCs. On the adaxial side, a thick SCW is deposited in a polar manner, with a characteristic composition of lignin, xylan, and cellulose synthesized by CESA7-containing CSCs. Within this polar SCW domain, lignin and cellulose are deposited concurrently, building up successive fine layers of lignified SCW material throughout stage 17 of fruit development. This mode of SCW formation contrasts with the sequence observed in many SCW-producing cell types, where lignification dominates later stages of SCW maturation and often coincides with programmed cell death (Meents et al., 2018).

The distinctive pattern of pectin methylesterification detected by LM20 in mature end*b* cells (Fig. 1A) points to a possible coordination between primary cell wall domains and SCW assembly. These pectin modifications are likely regulated by the activity of pectin methyl esterases (PME) and their inhibitors (PMEI) and have been reported in other contexts to provide anchoring platforms for cell wall remodeling enzymes or to facilitate direct interactions with lignin (although xylan plays the dominant role in mediating lignin interactions as we discuss below) (Francoz et al., 2019; Xiao et al., 2025). The reduced pectin content observed in *cesa7-1* valves, indicated by galacturonic acid measurements (Fig. 1D), may further reflect a potential coordination between primary and SCW processes. Investigating the mechanistic links between primary cell wall remodeling and end*b* SCW formation will be an important focus for future studies.

We found that *CESA7* was responsible for synthesizing more than half of the crystalline cellulose in *C. hirsuta* fruit valves, with the majority localized to end*b* SCWs. Despite the drastic depletion of cellulose in *cesa7* end*b* SCWs, thick lignin- and xylan-rich walls were still deposited on the adaxial side of the cells, and hinged regions formed that resembled the wild-type end*b* SCW pattern. These observations indicate that the initial spatial deposition of SCW polymers in *C. hirsuta* end*b* cells can occur independently of cellulose. Similar findings were described for the banded SCW pattern of protoxylem vessels in Arabidopsis *cesa7* mutants using an ectopic xylem induction system (Takenaka et al., 2018). In these cells, xylan and lignin were patterned in the absence of SCW cellulose in a microtubule-dependent manner (Takenaka et al., 2018). Together with our results, these data support a model in which the secretory pathway independently delivers CESAs, hemicelluloses and oxidative enzymes to microtubule-dense regions of the plasma membrane, where exocytosis establishes SCW patterns (Meents et al., 2018). Consistent with this model, xylan and lignin are deposited independently of cellulose in *C. hirsuta cesa7* end*b* SCWs, whereas cellulose is deposited independently of lignin in *lac4 11 17* mutants (Perez-Anton et al., 2022). Our results extend this framework by showing that although SCW patterning can be initiated without cellulose, maintaining these patterns requires SCW cellulose and likely relies on polymer maturation and interactions that have not yet been elucidated.

Cortical microtubules are essential in *C. hirsuta* end*b* cells for the spatiotemporal patterning of SCW domains along the adaxial cell edges, where all SCW polymers are precisely depleted to form thin hinges. However, the polar deposition of SCW polymers on the adaxial side of end*b* cells remained largely unchanged following microtubule disruption. Xylan deposition appeared unaffected, whereas a thin layer of ectopic lignin was observed throughout the end*b* cell wall (Fig. 7B), similar to that induced by increased *LAC11* gene expression (Perez-Anton et al., 2022). This relative independence of polar SCW patterning from microtubules in *C. hirsuta* end*b* cells contrasts with protoxylem cells, where microtubules are essential for establishing the banded SCW pattern (Takenaka et al., 2018). Therefore, identifying factors that specify the polarity of SCW deposition in *C. hirsuta* end*b* cells will be an important direction for future studies.

Our findings show that *CESA7* is essential to maintain the precise SCW patterns of end*b* and metaxylem cells in *C. hirsuta*. This requirement likely reflects the organization and physical interactions of cellulose, xylan, and lignin in the SCW, particularly the interactions between cellulose and xylan. Xylan adopts a flat ribbon (twofold) conformation when bound to cellulose (Simmons et al., 2016; Grantham et al., 2017). This conformation is induced by cellulose binding as xylan reverts to a non-flat (threefold) conformation in the Arabidopsis *cesa7* mutant *irx3-7* (Simmons et al., 2016). Lignin, in turn, preferentially binds non-flat conformations of xylan (Kang et al., 2019). Thus, the ability of xylan to adopt different conformations allows it to interact with both cellulose microfibrils and lignin, bridging the assembly of all three polymers in the SCW. Consistent with this, we found that the organization of *C. hirsuta* end*b* SCWs into flattened layers depended on *CESA7*-mediated cellulose synthesis and correlated with the ability to maintain highly ordered SCW patterns. Thus, cellulose-xylan interactions may scaffold polymer assembly to form a layered and precisely patterned SCW in end*b* cells. In the absence of SCW cellulose in *cesa7* end*b* cells, xylan and lignin failed to assemble into a layered structure, instead forming an irregular, undulating SCW that occupied a larger area (Figs. 4A-C, 5A-B). This overgrowth of *cesa7* end*b* SCWs was markedly enhanced by microtubule disruption, suggesting a synergistic interaction between *CESA7* and microtubules in SCW patterning. Together with previous findings from ectopic xylem induction systems (Takenaka et al., 2018; Pfaff et al., 2024), our results link the organization and interaction of SCW polymers to the formation of SCW patterns.

We found that cellulose deficiency altered the size and shape of pits in metaxylem cells in *C. hirsuta cesa7* roots. Similar pit defects have previously been reported in rice and Arabidopsis mutants deficient in xylan rather than cellulose (Wang et al., 2022). Xylan-rich domains were identified at pit borders, and their formation required xylan synthesis by IRX10 (Wang et al., 2022). Loss-of-function mutants had larger, irregular pits (Wang et al., 2022), resembling those observed in *C. hirsuta cesa7* mutants (Fig. 4D-F). In both cases, xylan-cellulose interactions appear critical for the compact organization and packing of SCW polymers to form smooth pit borders. Furthermore, a quantitative trait locus controlling pit size and shape in rice was recently identified as a xylan deacetylase encoded by *GDSL ESTERASE/LIPASE PROTEIN 33* (*GELP33*) (Zhang et al., 2025). An elite allele of *GELP33* modified xylans to a hypoacetylated state, enhancing xylan-cellulose binding at pit borders and thereby improving xylem hydraulic efficiency and crop yield in rice (Zhang et al., 2025). Key to these findings was resolving the three-dimensional structure of pitted SCWs at high resolution (Zhang et al., 2025). Extending similar analyses to SCWs in *C. hirsuta cesa7* mutants, along with investigating the specific role of xylan deposition in end*b* SCWs, represent promising avenues for future research into how SCW polymer assembly shapes specialized SCW patterns.

Our findings reveal both similarities and key differences between SCW patterning in end*b* cells and the well-studied process in xylem vessels. We propose that end*b* cells in the explosive fruit of *C. hirsuta* provide a valuable system to study SCW patterning, combining the advantages of a clear mechanical function in a non-essential cell type, with a novel cell polarity component absent from xylem cells. Comparative studies in this system are likely to uncover general principles of SCW patterning and the degree to which these mechanisms are conserved across diverse cell types with specialized functions in different plants. Moreover, biomechanical analyses integrating SCW ultrastructure with tissue mechanics could further reveal how the specific architecture of end*b* SCWs enables the storage and rapid release of elastic energy that powers explosive seed dispersal in *C. hirsuta*.

## Materials & Methods

### Plant materials and growth conditions

*Cardamine hirsuta* (Ox), herbarium specimen voucher Hay 1 (OXF) (Hay and Tsiantis, 2006) and Arabidopsis Col-0 were used as wild-type genotypes throughout this study. Plants grown on soil were cultivated in the greenhouse in long-day conditions (days: 20°C, 16 h; nights: 18°C, 8 h). Transgenic plants were generated by the floral dip method using *Agrobacterium tumefaciens* GV3101. *C. hirsuta p35S::GFP:TUA6* plants were described previously (Hofhuis et al., 2016). Arabidopsis *irx3-4* is a SALK T-DNA insertion line (SALK_029940) in Col-0 from NASC (N529940) and *cev1* is an EMS allele in Col-0 (Ellis et al., 2002) (gift from A. Sampathkumar).

### Plasmid construction and plant transformation

*pCESA7::mNG:CESA7* (*mNeonGreen*) and *pCESA7::3xGFP:NLS* plasmids were constructed by GreenGate cloning after all BsaI sites were mutagenized as described (Lampropoulos et al., 2013). The *CESA7* promoter (2000bp before ATG) was PCR-amplified from *C. hirsuta* genomic DNA and cloned into pGGA000 entry clone. The *CESA7* cDNA (3114bp including STOP) was PCR-amplified from *C. hirsuta* cDNA and cloned into pGGC000 entry clone. The *mNeonGreen* sequence was amplified from *CESA7pro::mNG-CESA7* (gift from L. Samuels) and cloned into pGGB entry clone. Entry vector combinations were cloned into the pGGZwf01 binary vector (Perez-Anton et al., 2022) as described (Table S2 for primers and GG modules). *pCESA7::3xGFP:NLS* was transformed into wild-type *C. hirsuta* and 9 independent T_2_ lines were analyzed. *pCESA7::mNG:CESA7* was transformed into Arabidopsis *irx3-4* and *C. hirsuta cesa7-1* homozygous mutants. Complementation (restoration of plant height, fruit length, end*b* SCW patterning) was observed in 27 of 30 independent T_2_ lines of *C. hirsuta cesa7-1* and 67 of 70 independent T_2_ lines of Arabidopsis *irx3-4*.

The *pUBQ10::GR-LhG4/pOp6::PHS1ΔP:mCherry* plasmid was constructed as a multiple expression GreenGate cassette by combining two previously described modules into the pGGZwf01 binary vector (Lampropoulos et al., 2013; Schurholz et al., 2018; Vilches Barro et al., 2019; Perez-Anton et al., 2022). The final construct was transformed into *C. hirsuta p35S::GFP:TUA6* plants and 7 of 29 independent T_2_ lines showed strong dexamethasone induction of PHS1ΔP:mCherry.

### CRISPR/Cas9 mutagenesis of *CESA7*

CRISPR/Cas9 directed mutagenesis of the *C. hirsuta CESA7* gene was performed using MultiSite Gateway cloning as previously described (Alvim Kamei et al., 2020; Perez-Anton et al., 2022). The first entry vector contained four *Streptococcus pyogenes* (Sp) Cas9-compatible sgRNA sequences targeting *CESA7* that were identified and evaluated based on location, high efficiency and minimum off-target prediction using CCTop (Stemmer et al., 2015) (Table S1, Fig. S2). This vector containing the four sgRNA sequences each driven by the Arabidopsis U6 RNA pol III promoter was synthesized by GeneScript into a single Gateway-compatible entry vector with attL2-L5 sites. The second entry vector contained the S*pCas9 sequence* driven by an egg-cell specific promoter (EC1.2en-EC1.1p) with attL1-R5 sites. A LR-reaction was performed to combine the two entry clones with a Gateway-compatible pPZP200-based binary vector (pPZP200-FAST-RFP) with attR1-R2 sites. This construct was transformed into *C. hirsuta* wild-type plants. Positive transformants were selected based on the presence of seed fluorescence and genotyped with primers that amplified regions targeted by the four sgRNAs (Table S2). Non-fluorescent T_2_ seeds were selected from independent lines with *CESA7* mutations and genotyped to find plants with homozygous mutations. Three different alleles, *cesa7-1*, *cesa7-2* and *cesa7-3*, were used in this study (Fig. S2).

### Photography

Photographs of plants, fruit and fruit valves were taken with a Nikon D800 equipped with either an AF-S Micro NIKKOR 105 mm 1 : 2.8 G ED or AF-S NIKKOR 24–85 mm 1 : 3.5–4.5 G objectives.

### Microscopy

To prepare fruit cross-sections, whole fruits were embedded in 1.5 ml tubes containing 5% low melting agarose (Hi-Pure Low agarose; Biogene Ltd) and cut into 100 μm sections using a Leica Vibratome VT1000 S. For intact fruit valves, mature valves were peeled from the fruit. Samples were fixed, cleared, and stained using an adapted Clearsee protocol as previously described (Ursache et al., 2018; Perez-Anton et al., 2022). To stain cellulose, sections were stained with either 0.1% direct red 23 (Sigma-Aldrich) for two hours or 0.1% calcofluor white (Sigma-Aldrich) for 16 hours. Lignin was stained with 0.2% basic fuchsin (Sigma-Aldrich) for 16 hours. To image *pCESA7::3xGFP:NLS* in root samples, seedlings were stained with 10 μg/ml propidium iodide for 5 min before imaging.

A Leica TCS SP8 was used for confocal laser scanning microscopy (CLSM) with either a HCX PL APO lambda blue (63x/1.20 water), a HC FLUOTAR L (25x/0.95 water), or a HC PL FLUOTAR (10x/0.30 dry) objective lens. 0.3-0.8 μm z-stack slices were acquired. The excitation and detection windows were set as follows: lignin autofluorescence (405 nm, 410-545 nm), direct red 23 (561 nm, 560-650 nm), calcofluor white (405 nm, 425-475 nm), basic fuchsin (561nm, 600-655 nm), GFP (488 nm, 500-540 nm) and PI (488 nm, 600-690 nm). For experiments comparing fluorescence intensities of the stains, image acquisition settings were kept the same.

A Leica SP8 FALCON-DIVE multiphoton CLSM was used to image the end*b* SCW of intact fruit valves. Fixed and cleared fruit valves were sealed in between a glass slide and coverslip, with the endocarp facing the coverslip, and imaged with a HC PL APO CS2 (63x/1.3 glycerol) objective lens. To visualize the lignin, the laser was set to an 825 nm wavelength and a 440-500 emission bandpass filter. 0.2-0.3 μm z-stack slices were acquired of the entire depth of the SCW.

A Leica SP8 FALCON-DIVE multiphoton CLSM was also used to image microtubules and the end*b* SCW in *C. hirsuta p35S::GFP:TUA6* fruits. Valves were peeled from early stage 17b fruit and sliced into 2 mm sections to maintain end*b* cell viability. The sections were then carefully mounted in perfluoroperhydrophenanthrene (Sigma-Aldrich) between a coverslip and glass slide with the endocarp facing the coverslip. After sealing the coverslip to the glass slide, samples rested for ten minutes to avoid sample drift during imaging, and were then imaged with a HC PL APO CS2 (63x/1.3 glycerol) objective lens. Excitation and detection windows (nm) and laser power (%) were set as follows: lignin autofluorescence (825 nm, 440-500 nm, 4%), GFP (930 nm, 490-550 nm, 3%). 0.2 μm z-stack slices were acquired of the entire depth of the SCW before switching channels.

### Image analysis

ImageJ was used to quantify the area and fluorescence intensity of the end*b* SCW of fruit sections in C*. hirsuta* and Arabidopsis. Regions of interest (ROI) were first selected using the wand tracing tool on the lignin image and saved in the ROI manager. The ROIs were then superimposed on the cellulose image. Area, Integrated Density and Mean Grey Value for each ROI were then measured. Five ROIs of the background were then selected and the mean fluorescence were also measured. To quantify the fluorescence intensity for each ROI, the following equation was used: corrected total cell fluorescence (CTCF) = Integrated Density – (Area of Selected Cell x Mean Grey Value of background readings).

To analyze the end*b* SCW in intact fruit valves, maximum-intensity projections of the z-stacks were done in ImageJ. Three-dimensional renderings of the end*b* SCWs and microtubules were done with Imaris software. Microtubule confocal images were first processed in ImageJ: Background Subtraction (50 rolling ball radius), Enhance Contrast, Smooth. Both microtubule and lignin channels were uploaded into the Imaris software as separate z-stacks with the volume rendering set to Maximum-Intensity Projections (MIP). Depending on the desired visualization, custom clipping planes were set and the images were rotated to specific angle.

For multivariate shape analysis using the LeafI software (Zhang et al., 2020), lignin confocal images were transformed into binary images using ImageJ. To create the binary image, the outline of the cell wall was selected using the Magic Wand tool and the outline was then “filled” using the ROI manager. All images were created so that they had the same orientation, dimensions, and pixel size. For the LeafI settings, the landmarks of the SCW were manually defined as the midpoint of the top and bottom of the adaxial SCW. Registration was defined as “Align main-axis (2-point)”. For shape space, only the “normalize” setting was selected.

To analyze the root metaxylem SCW, wild-type and *cesa7-1* seeds were sown onto 0.5 MS-sucrose plates and stratified in the dark at 4°C for three days. Plants were grown in long-day conditions for six days before fixing, clearing and staining the seedlings as previously described (Ursache et al., 2018). After confocal imaging, the ellipse function of ImageJ was used on the confocal images to measure the area of the metaxylem pits.

### Immunofluorescence labelling of fruit sections

Early stage 17b fruits were processed by high-pressure freezing, freeze substitution and resin embedding as previously described (Neumann and Hay, 2020). Immunofluorescence labelling of different cell wall epitopes on semithin (1 µm) sections was performed according to (Neumann and Hay, 2020), using the following primary, rat monoclonal antibodies (diluted 1 in 10): LM11 (heteroxylans), LM19, LM20 (pectins) and LM25 (xyloglucans). The initial blocking step was carried out at room temperature for 30 minutes. Prior to the detection of xyloglucans, sections were subjected to digestion by pectinase (Sigma-Aldrich Germany, 17389) or macerozyme (Duchefa M8002) respectively at 1U/mL, adopted from (Wilson et al., 2015). Images were acquired with either a Zeiss Axiovert fluorescence microscope or a Leica TCS SP8 CLSM.

### Transmission electron microscopy

Early stage 17b fruits were processed by high-pressure freezing and freeze substitution, embedded into LR White resin, sectioned and then contrasted according to (Neumann and Hay, 2020). Micrographs were taken with a Hitachi HT-7800 TEM (Hitachi High-Technologies Europe GmbH, Krefeld, Germany) operating at 100 kV fitted with an EMSIS Xarosa digital camera.

### Cryo-fracture scanning electron microscopy

Fresh stage 17b fruit of *C. hirsuta* wild type and *cesa7-1* were flash-frozen with liquid nitrogen and subjected to cryo-fracture, sublimation and gold-coating in an Emitech K1250x cryo unit. Images were taken using a Zeiss Supra 40VP scanning electron microscope operating at 3 kV.

### Chemical treatments

#### Dexamethasone (Dex) induction of PHS1ΔP mCherry

7 mm (stage 15) fruits (still attached to the petiole and a portion of the stem) were selected from *pUBQ10::GR-LhG4/pOp6::PHS1ΔP:mCherry; p35S::GFP:TUA6* plants and placed into 190 mL polystyrene plant tissue culture chambers (Greiner Bio-One) filled with 0.5 MS agar media supplemented with 2% sucrose and 0.1% Plant Preservative Mixture (Plant Cell Technology). For the Dex treatment, a final concentration of 1 mM Dex (Sigma-Aldrich) dissolved in water was used. The chambers were transferred to a growth room in long day (16-hour light) conditions. After five days in the chambers, microtubule depolymerization in the end*b* was checked by imaging the cells from a peeled fruit valve with a Leica SP8 CLSM. Fruits were sectioned, fixed, and cleared after 11 days of growth in the chamber. Alternatively, 7 mm fruit were treated daily by dipping in water or 2 mM Dex solutions with 0.02% Silwet L-77 surfactant added, and valve coiling was assessed in exploded fruit after 10 days.

#### Oryzalin treatment

Stage 16 fruits (10-13 mm in length) from wild-type, *cesa7-1* and *35S::GFP:TUA6* plants were dipped in a solution containing 1 mM oryzalin (Sigma-Aldrich), 5.8% DMSO, and 0.02% Silwett for 30 seconds. Fruits were dipped every day for nine days before sectioning, fixing, and clearing. To confirm that the treatment was sufficient to induce microtubule depolymerization, *35S::GFP:TUA6* fruits were screened with a Leica SP8 CLSM. *Isoxaben and DCB treatments:* For isoxaben treatments, 7 mm (stage 15) or 19-20 mm (stage 17a) fruits (still attached to the petiole and a portion of the stem) were carefully placed into 190 mL polystyrene plant tissue culture chambers (Greiner Bio-One) filled with 0.5 MS agar media supplemented with 2% sucrose and 0.1% Plant Preservative Mixture (Plant Cell Technology). A final concentration of 40 μM isoxaben (Supelco) dissolved in DMSO or DMSO alone (for the mock treatment) was added to the media. The chambers were transferred to a growth room in long day (16-hour light) conditions. Fruits were sectioned, fixed, and cleared after eight days of growth in the chamber. MorphograpX (Barbier de Reuille et al., 2015) was used to measure the exocarp cell area from confocal micrographs of fruit cross-sections. For DCB treatments, 19-20 mm (stage 17a) fruits were grown in the same conditions as the isoxaben assay except supplemented with 10 μM DCB (Sigma-Aldrich) dissolved in DMSO or DMSO alone (for the mock treatment). Fruits were sectioned, fixed, and cleared after eight days of growth.

### Cell wall analysis

Fruit valves were collected from mature fruits (approximately 530 valves for wild type and 660 valves for *cesa7-1*) and flash frozen in liquid nitrogen before being stored at 80°C until analysis. Samples were freeze-dried and aliquoted into three technical replicates per genotype (approximately 90 mg dry weight for both wild type and *cesa7-1*). Cellulose content and matrix polysaccharide-derived monosaccharide composition including uronic acids of the samples were determined as described (Bauer and Ibanez, 2014; Yeats et al., 2016). In short, Alcohol Insoluble Residue (AIR) was prepared from freeze-dried samples and split into two samples. One half of the samples were treated with a weak acid (4% sulfuric acid) to release matrix polysaccharide-derived monosaccharides, while the other half of the samples were treated initially with a strong acid (72% sulfuric acid) followed by diluting the sulfuric acid concentration to 4% to yield monosaccharides both derived from cellulose and the matrix polymers. Subtraction of the two values allows for the quantification of crystalline cellulose. Monosaccharides of all fractions were quantified using an IC Vario high-performance anion-exchange chromatography system 1068 (Metrohm, Herisau, Switzerland) equipped with a CarboPac PA20 column (Thermo Fisher Scientific, Waltham, MA, USA) and an amperometric detector (Metrohm) using a sodium hydroxide gradient. Acetyl-bromide lignin content of the AIR samples was determined as previously described (Foster et al., 2010).

### RNA sequencing and analysis

Valves from stage 17ab fruits (16 valves from eight fruits per replicate) were flash frozen in liquid nitrogen and RNA was extracted from three biological replicates of *C. hirsuta* wild type and *cesa7-1* using Spectrum^TM^ Plant Total RNA kit (Sigma-Aldrich). For Illumina sequencing, the cDNA synthesis, Stranded Poly-A selection library preparation (two-sided, 150bp) and sequencing were carried out by Novogene using the Illumina NovaSeq6000 platform. Data was analyzed using a local Galaxy instance (Galaxy, 2024). Principal component analysis showed that one *cesa7-1* replicate did not cluster with the other replicates and was discarded from further analysis. RNA quality was checked using FastQC, reads were mapped to the *C. hirsuta* reference genome (Gan et al., 2016) using RNA STAR, raw read counts were quantified with featureCounts, and differentially expressed genes (DEGs) were identified using DESeq2. DEGs were selected based on a cut-off with a fold-change (FC) of 2, and a p-adjusted value of 0.05. For gene ontology (GO) analysis, we used *A. thaliana* GO term annotations for orthologous *C. hirsuta* genes (Gan et al., 2016). Analyses were carried out in R v.4.2.1 using the package ClusterProfiler v.4.4.4 for GO analysis (Wu et al., 2021) and ggplot2 v.3.5.2 for volcano plots (Wickham, 2016).

### Statistical analysis

Statistical analyses were done with R v4.2.1 (R Core Team, 2024).

### Accession numbers

Short read sequence data from this article has been deposited in the European Nucleotide Archive (ENA) at the European Molecular Biology Laboratory’s European Bioinformatics Institute (EMBL-EBI) under accession number PRJEB76589 (https://www.ebi.ac.uk/ena/browser/view/PRJEB76589). Accession numbers for major genes reported in *C. hirsuta*: CARHR195790 (*CESA7*) and discussed in Arabidopsis: AT5G17420 (*CESA7*), AT5G05170 (*CESA3*).

## Supplemental data

Figure S1. Arabidopsis *CESA7* allele *irx3-4* lacks cellulose in end*b* SCWs, supports Fig. 2.

Figure S2. CRISPR/Cas9 alleles of *C. hirsuta CESA7*, supports Fig. 2.

Figure S3. *C. hirsuta CESA7* controls SCW cellulose synthesis, supports Figs. 2 and 3.

Figure S4. Cellulose synthesis inhibitors DCB (2,6-dichlorobenzonitrile) and isoxaben had no effect on end*b* SCWs in *C. hirsuta* fruit, supports Fig. 3.

Figure S5. Immunofluorescence detection of xylan in *C. hirsuta* fruit end*b* SCW cross sections using LM11 antibody, supports Figs. 2 and 7.

Figure S6. Effects of oryzalin treatment on wild-type and *cesa7* end*b* SCWs, supports Fig. 8.

Table S1. RNAseq analysis of wild-type vs *cesa7-1* fruit valves.

Table S2. Primers, sgRNAs and GreenGate modules.

## Acknowledgements

We thank L. Samuels and A. Sampathkumar for sharing materials, K. Lufen for cell wall analyses, W. Faigl for technical support, S. Dilip Pophaly for bioinformatics support and R. Franzen for scanning electron microscopy. This work was supported by the Deutsche Forschungsgemeinschaft (DFG) under Germany’s Excellence Strategy—EXC 2048/1—Project ID: 390686111 to M.P. and DFG FOR2581 Plant Morphodynamics grant to A.H.

## Author contributions

Conceptualization, A.H. and R.C.E.; Investigation, R.C.E., U.N. and A.E; Cell wall analysis, M.P.; Writing, A.H. and R.C.E; Funding Acquisition, A.H. and M.P.; Supervision, A.H.

