## Supplemental Figures S1-S6 for "CESA7 and microtubules pattern complex secondary cell walls in explosive fruit"

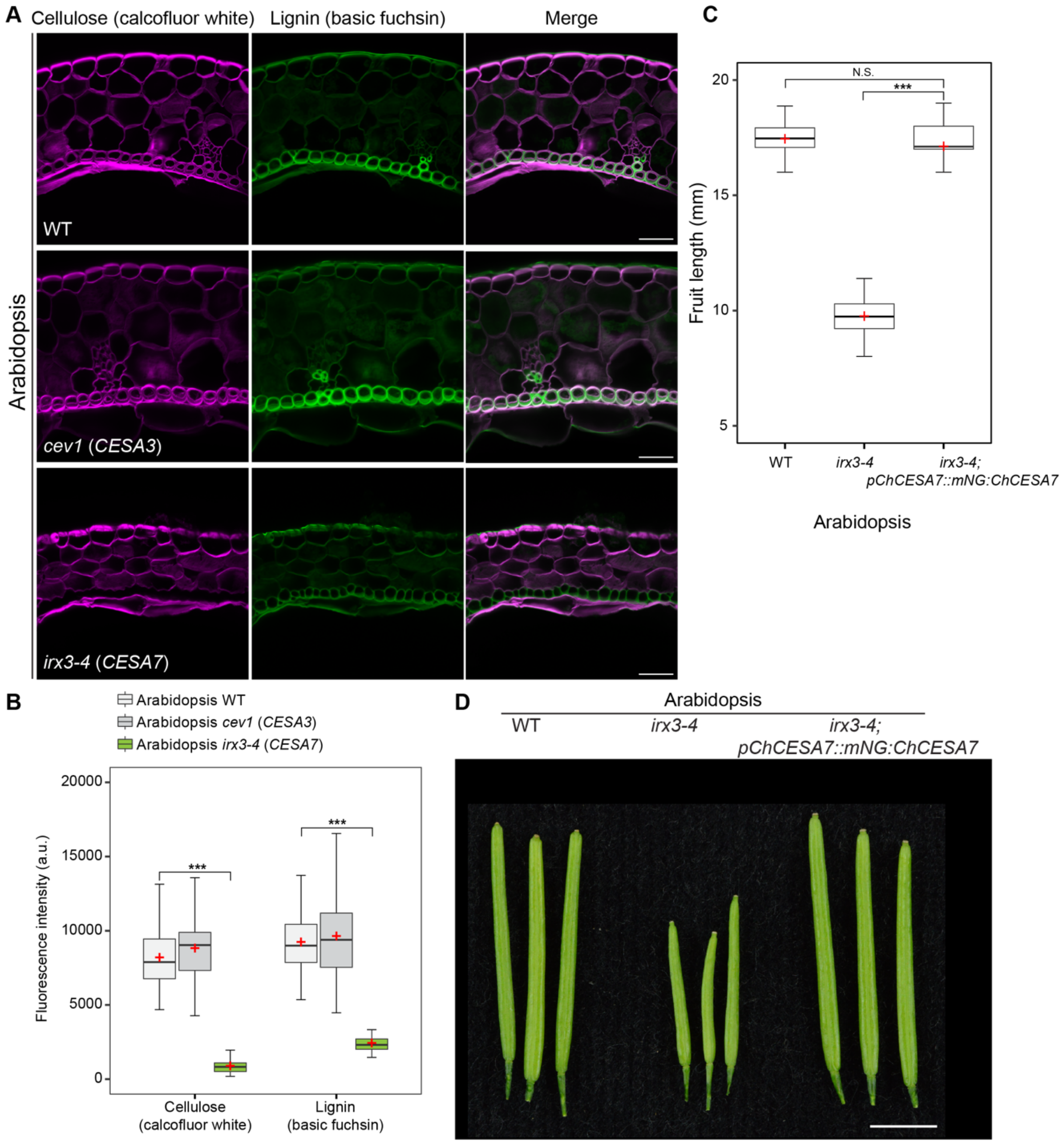


**Figure S1. Arabidopsis *CESA7* allele *irx3-4* lacks cellulose in end*b* SCWs.** Supports Figure 2. **(A)** CLSM of Arabidopsis stage 17b fruit valves showing cellulose stained with calcofluor white (magenta), lignin stained with basic fuchsin (green) and both channels merged in wild type, *cev1* (*CESA3* allele) and *irx3-4* (*CESA7* allele). **(B)** Boxplot of fluorescence intensity, measured in arbitrary units (a.u.), of calcofluor white and basic fuchsin signals in end*b* SCWs of Arabidopsis wild-type (n = 70), *cev1* (n = 75) and *irx3-4* (n = 70) fruit valves shown in (A). **(C)** Boxplot of fruit length in Arabidopsis wild-type, *irx3-4* and *irx3-4* complemented with *C. hirsuta pCESA7::mNG:CESA7* (n = 40 per genotype). **(D)** Fruits of Arabidopsis wild type, *irx3-4* and *irx3-4* complemented with *C. hirsuta pCESA7::mNG:CESA7*. Plots in (B-C) show median (thick black line) and mean (red cross). *** denotes statistical significance at $P < 0.001$ using Wilcoxon rank sum test. N.S. = not statistically significant with $P > 0.05$ using Wilcoxon rank sum test. Scale bars: 30 µm (A), 5 mm.

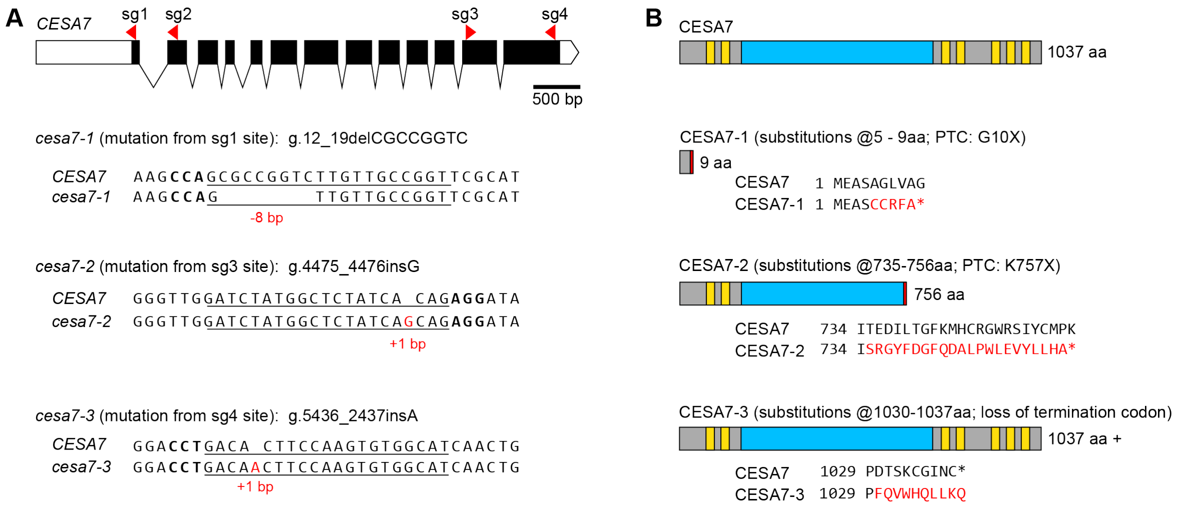


**Figure S2. CRISPR/Cas9 alleles of *C. hirsuta CESA7*.** Supports Figure 2. **(A)** Gene model of *C. hirsuta CESA7*; details described in legend; scale bar: 500 bp. Description of *cesa7-1*, *cesa7-2* and *cesa7-3* mutant alleles. Each mutation is indicated in red and its position counting from the start codon in genomic sequence (g.). For example, g12_19del indicates a deletion of eight nucleotides. Legend: UTRs (white bars), exons (black bars), introns (lines), sgRNA location and direction (red arrowheads), PAM sequence (bold), sgRNA sequence (underline) and mutation (red). **(B)** Schematic of wild-type *C. hirsuta* CESA7 protein and putative translational products of *cesa7-1*, *cesa7-2* and *cesa7-3* mutant alleles indicating the number of amino acid (aa) residues resulting from missense translation and PTC; details described in legend. For example, "substitutions@5-9aa; PTC: G10X" indicates that aa residues 5 to 9 result from missense translation, followed by a PTC at G10. Legend: transmembrane domains (yellow bars), catalytic domain (blue bar), premature termination codon (PTC, red bar), termination codon (*) and altered amino acid (aa) sequence (red).

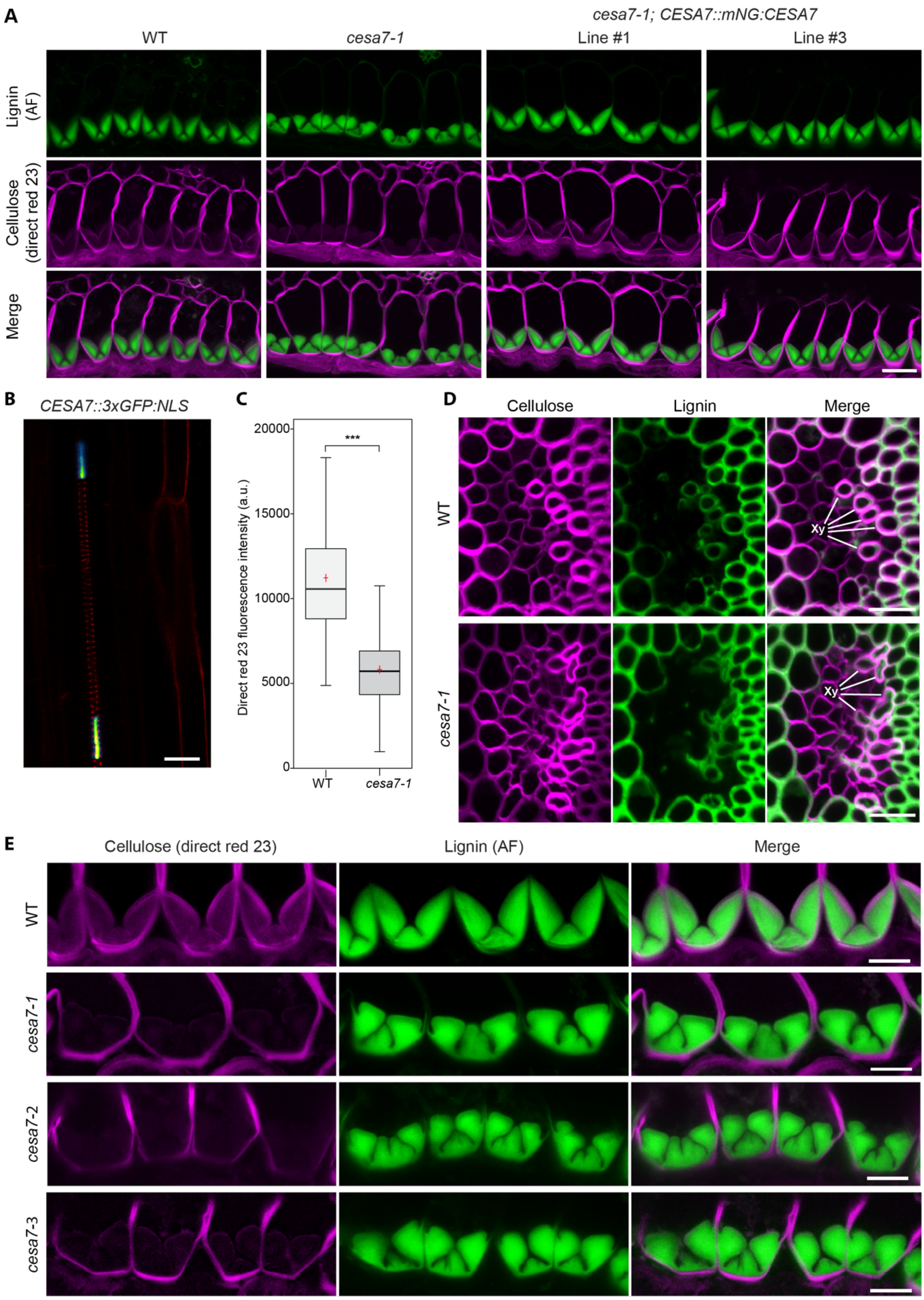


**Figure S3. *C. hirsuta CESA7* controls SCW cellulose synthesis.** Supports Figures 2 and 3. **(A)** CLSM of *C. hirsuta* end*b* SCWs in stage 17b fruit of wild type, *cesa7-1* and two *cesa7-1* independent transgenic lines complemented with *C. hirsuta pCESA7::mNG:CESA7* showing lignin autofluorescence (green), cellulose stained with direct red 23 (magenta) and both channels merged. **(B)** Expression of *C. hirsuta pCESA7::3×GFP:NLS* (viridis LUT) in protoxylem cells, outlined in red with propidium iodide staining, in roots of 6-day old *C. hirsuta* seedlings. **(C)** Boxplot of direct red 23 fluorescence intensity, measured in arbitrary units (a.u.), in end*b* SCWs of *C. hirsuta* wild type (n = 75 cells from 5 fruits) and *cesa7-1* (n = 83 cells from 5 fruits) shown in Fig. 3A. Plots show median (thick black line) and mean (red cross). *** denotes statistical significance at $P < 0.001$ using Wilcoxon rank sum test. **(D)** CLSM of collapsed xylem vessels (Xy) in the replum of *C. hirsuta cesa7-1* fruit compared to wild-type showing cellulose, lignin and both channels merged. **(E)** *C. hirsuta* end*b* SCWs in stage 17b fruit of wild type, *cesa7-1*, *cesa7-2* and *cesa7-3* showing cellulose stained with direct red 23 (magenta), lignin autofluorescence (green) and both channels merged. Scale bars: 20 µm (A, D), 10 µm (B, E).

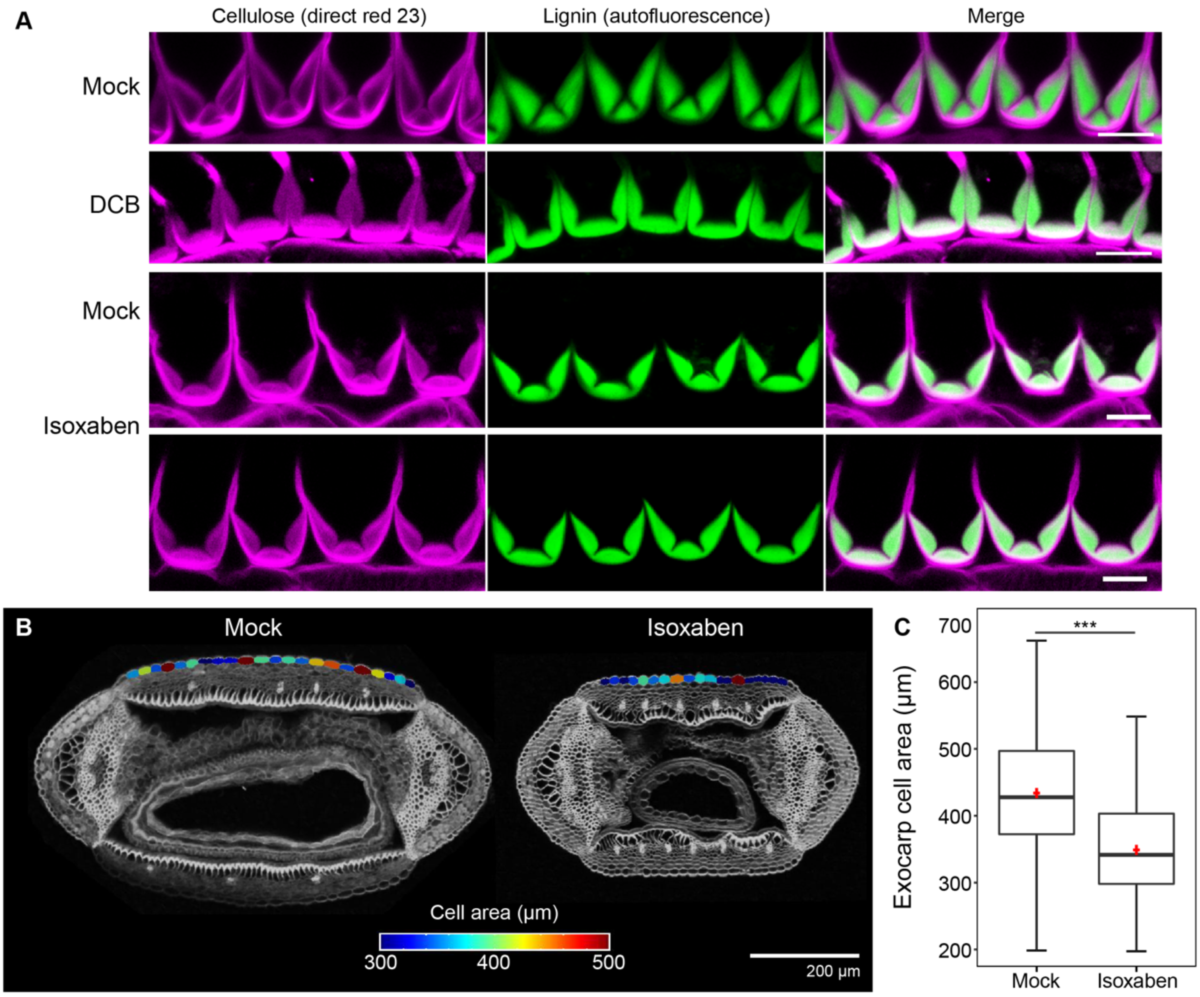


**Figure S4. Cellulose synthesis inhibitors DCB (2,6-dichlorobenzonitrile) and isoxaben had no effect on end*b* SCWs in *C. hirsuta* fruit.** Supports Figure 3. **(A)** CLSM of end*b* SCWs in stage 17b fruit of wild-type *C. hirsuta* treated with DCB or isoxaben, compared to mock treatments, showing cellulose stained with direct red 23 (magenta), lignin autofluorescence (green) and both channels merged. **(B)** Cross sections of stage 17b fruit of wild-type *C. hirsuta* treated with isoxaben, compared to mock treatment; heatmap indicates cell area in exocarp cells segmented with MorphoGraphX software. **(C)** Boxplot of exocarp cell area measured in 5 mock-treated valves (n = 106 cells) and 5 isoxaben-treated valves (n = 109 cells). Plot shows median (thick black line) and mean (red cross). *** denotes statistical significance using two-sample t-test with equal variance, $P < 0.001$. Scale bars: 10 µm (A), 200 µm (B).

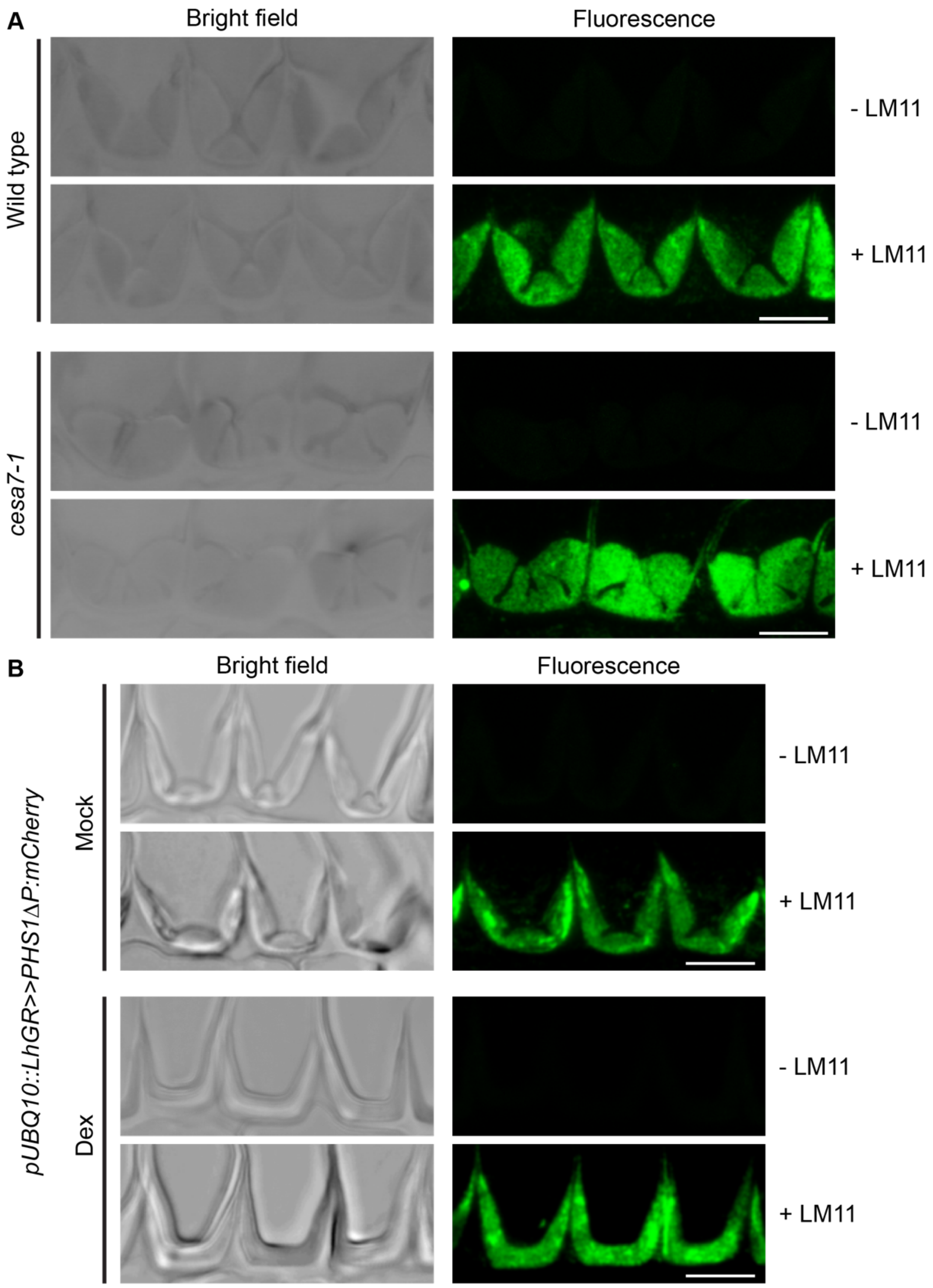


**Figure S5. Immunofluorescence detection of xylan in resin-embedded *C. hirsuta* fruit end*b* SCW cross sections using LM11 antibody.** Supports Figures 2 and 7. Brightfield and fluorescence micrographs of samples processed including (+LM11) or excluding (-LM11) primary LM11 antibody. **(A)** Wild-type and *cesa7-1* stage 17b fruit. Fluorescence panels also shown in Fig. 3A. **(B)** *pUBQ10::GR-LhG4/pOp6::PHS1ΔP:mCherry* fruit grown for 11 days on media without (mock) or with 1 mM dexamethasone (Dex). Fluorescence panels also shown in Fig. 7C. Scale bars: 10 µm.

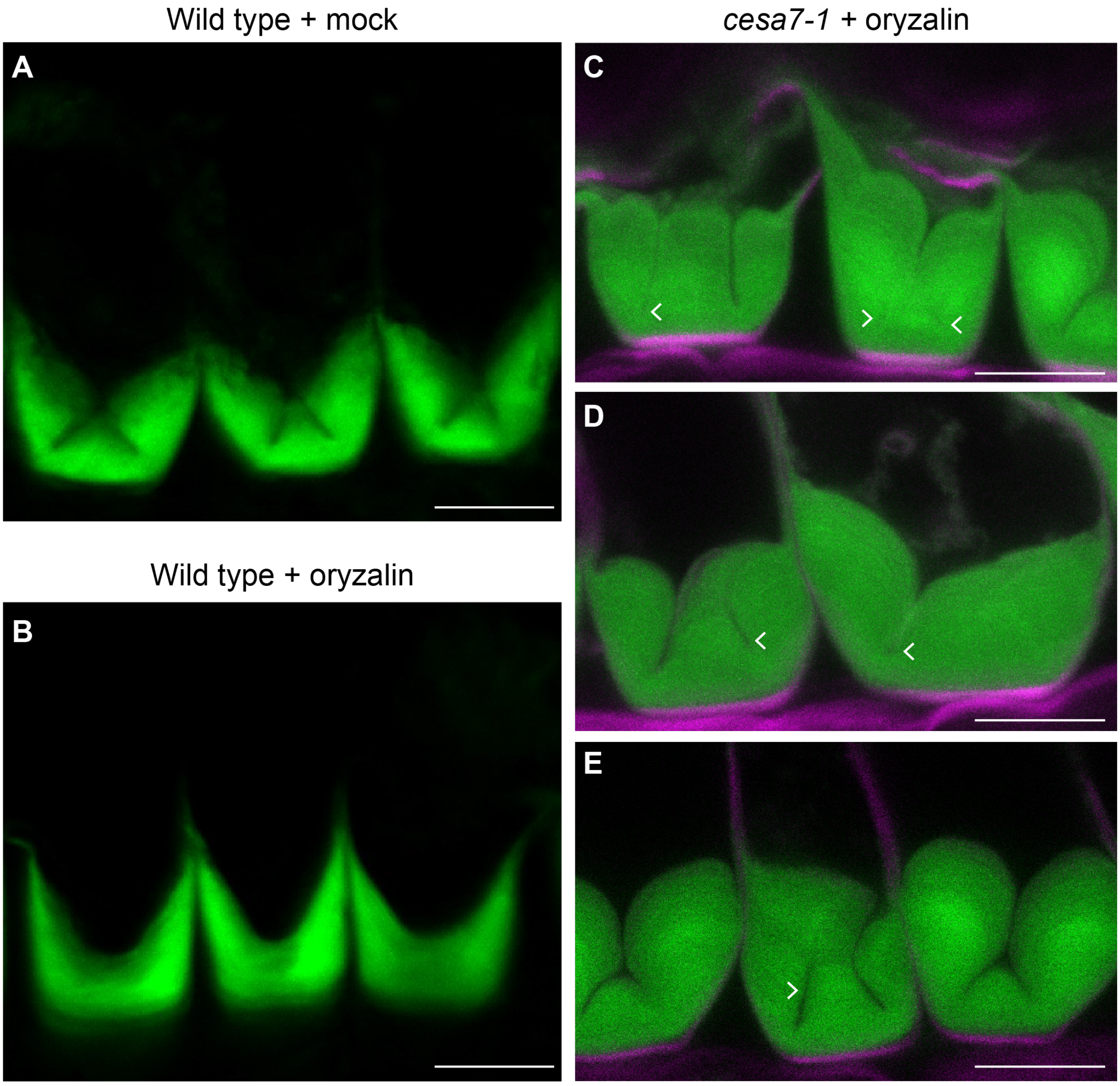


**Figure S6. Effects of oryzalin treatment on wild-type and *cesa7* end*b* SCWs.** Supports Figure 8. CLSM of end*b* SCWs in stage 17b fruit showing lignin autofluorescence (green, A-E) together with cellulose stained with direct red 23 (magenta, C-E). **(A-B)** Wild-type *C. hirsuta* after mock (A) or oryzalin (B) treatment. **(C-E)** *C. hirsuta cesa7-1* treated with oryzalin. Arrowheads indicate holes or gaps within the SCW where hinges were initiated, but not maintained, and subsequently overgrown by new SCW layers. Scale bars: 10 µm.
